# Folding of the cerebellar cortex is clade-specific in form, but universal in degree

**DOI:** 10.1101/2023.05.17.541232

**Authors:** Annaleigh R. York, Chet C. Sherwood, Paul R. Manger, Jon H. Kaas, Bruno Mota, Suzana Herculano-Houzel

## Abstract

Like the cerebral cortex, the surface of the cerebellum is repeatedly folded. Unlike the cerebral cortex, however, cerebellar folds in a given brain are much thinner and more numerous; repeat themselves largely along a single direction, forming long strips transverse to the mid-sagittal plane, like an accordion; and occur in the smallest of cerebella, including those of lissencephalic mammals and non-mammal vertebrates. We have shown previously that while the location of folds in mammalian cerebral cortex is clade-specific, the overall degree of folding strictly follows a universal power law relating cortical thickness, and the exposed and total surface areas. This law is derived from a statistical-physics model for gyrification that postulates that folding results from the interplay between axonal elongation dynamics and the self-avoiding nature of the expanding cortical surfaces. Since both aspects are present in the cerebellum, we hypothesize that a similar relation across species also exists therein. Furthermore, given the modular organization of cerebellar architecture and circuitry, as well as the transverse orientation of the folia, it is plausible that this relation is reflected in the degree of folding of the mid-sagittal section of the cerebellum, which greatly facilitates analysis. Here we show that a strict universal scaling law does apply to the folding of the mid-sagittal sections of the cerebellum of 53 species belonging to six mammalian clades, spanning a large range of sizes and degrees of gyrification. This folding is hierarchical and can be explicitly separated into branching orders, such that position of the 1^st^-order folds is largely stereotypical across all mammals examined. Subsequent nth-order folds become progressively less stereotypical, and folding within such cerebellar subsections scales with power laws whose exponents decrease monotonically with branching order, converging to the exponents predicted by a two-dimensional version of the same gyrification model that describes cortical folding. We propose that the changes in scaling exponent with branching order occurs as increasing amounts of white matter are included in the folding volume of the cerebellum, reflecting the difference between the outside-in development of the cerebellar cortex around a preexisting core of already connected white matter, compared to the inside-out development of the cerebral cortex with a white matter volume that develops as the cerebral cortex itself gains neurons. Our data strongly indicate that the mammalian cerebellum folds as a multi-fractal object, emerging from the interplay between clade-specificity and universality, and between phylogenetical contingency and the physics of self-organization. Thus, repeated folding, one of the most recognizable features of biology, can arise simply from the universal applicability of physical principles, without the need for invoking selective pressures in evolution; and diversity arises within the constraints imposed by physics.

## INTRODUCTION

Over 100 years ago, Ramón y Cajal established that the histological structure of the cerebellum, while distinct from that of the cerebral cortex, is nearly identical across all vertebrates examined (Ramón y Cajal, 1911), and so invariant that Cajal referred to it as a ‘law of biology.’ Similarly, the literature on cerebellar morphology has assumed that there is invariance also on the gross morphological level, in the form of a “common plan” of folding into ten lobules varying solely in size and degree of folding across species and clades (Sultan and Braitenberg, 1993; Sultan & Glickstein, 2007; Voogd & Glickstein, 1998). The concept of a common plan of folding of the cerebellum dates back to Bert Stroud’s conclusion that “there is a fundamental plan after which the cerebella of all mammals are formed” (Stroud, 1895), which implies that a human cerebellum is simply a “scaled-up” version of a mouse cerebellum in its morphology – even though Stroud’s conclusion was reached after studying the cerebella of only cats and humans.

The cerebellum does vary widely in absolute and relative volumes of grey and white matter across mammalian species (Bush & Allman, 2003; Maseko et al., 2012). While it has been noted that there is also variation in the relative abundance of folia and sizing of subdivisions (lobules) across species (Sultan & Glickstein, 1993), these discrepancies have sometimes been attributed to functional differences in characteristic behaviors, such as the elaborate climbing skills of squirrels (Sultan and Braitenberg, 1993; Sultan & Glickstein, 2007).

During our previous comparative studies of the brains of highly diverse mammalian species (Herculano-Houzel et al., 2006, 2007, 2011, 2014; Kazu et al., 2014; Neves et al., 2014; Jardim-Messeder et al., 2018; Dos Santos et al., 2018), we have not only observed variation in the morphological appearance of the cerebellum belonging to different species, but interestingly that the shape of the midsagittal surface is distinctive to the point that cerebella can be identified as belonging to rodent, marsupial, carnivoran, artiodactyl or primate species by simple visual inspection. Such a readily apparent clade-distinctive morphology has long been known in the cerebral cortex (Welker, 1990), but is yet to be shown in the cerebellum.

Despite the clade-specific spatial patterns of cerebral cortical folding, we have shown recently that the degree of folding, captured by the relationship between exposed cortical surface area and total cortical surface area, is fully predictable from physical principles according to a single power law that describes the conformation of minimal effective free energy, that is, the most energetically stable and therefore likely conformation, of self-avoiding expanding surfaces like the cerebral cortex (Mota & Herculano-Houzel, 2015). This power law describes how the larger the surface area of a cerebral cortex of constant thickness, or the thinner the cortex of a similar surface area across species, the more it folds – regardless of the number of cells that compose the structure (Mota and Herculano-Houzel, 2015). This power law also describes individual cerebral cortices across length scales, indicating that they are well-approximated by a fractal of dimension 2.5 (Losa, 2011; Kaandorp, 1994; Wang, 2022). Fundamentally, it can be shown from first principles that the degree of folding is simply a function of the total area expressed as a multiple of the squared cortical thickness, that is, folding scales not with surface area but with the ratio between surface area and squared cortical thickness (Mota and Herculano-Houzel, 2015). This prediction corroborates previous qualitative demonstrations that, across mammalian species, thinner cortices are more folded (Pillay and Manger, 2007; Manger et al., 2012). If such a mechanism applies universally to determine the degree of folding of a layered brain structure that expands in development, then it should capture equally well the diversity in degree of folding of the cerebellar surface across species. In this case, all else being equal, a much thinner and broader sheet like the cerebellar cortex would be expected to have much more of its surface tucked into folds than the cerebral cortex, as is the case.

Here we test the hypothesis that the same physical principles that describe the degree of folding of the cerebral cortex also apply to the cerebellum, even though the two structures originate from different anlage in the neural tube: the cerebral cortex from the dorsal ventricular zone of the second neuromere, and the cerebellum from the isthmus and first hindbrain neuromeres (Watson, Mitchelle, & Puelles, 2016). Importantly, the pattern of neurogenesis also differs between the two cortices. In the cerebral cortex, neurogenesis occurs in an inside-out manner, with neurons mostly originating from the sub-ventricular zone and radiating *outwards* as they form simultaneously grey matter and subcortical white matter. In contrast, the cerebellar cortex develops in two phases: first inside-out, as the deep nuclei and the Purkinje cells are born and connect to each other, forming the white matter core of the cerebellum; and then outside-in, as the external cerebellar germinal layer gives rise to large numbers of granule cell neurons that migrate inwards, past the preexisting Purkinje cells, and lead to the formation of lobules that expand massively in surface area, much more than they gain in thickness, on top of the preexisting white matter (Goldowitz and Hamre, 1998). This differential expansion of the outer layer of the cerebellar cortex was recently shown to initiate the placement of cerebellar folds, followed by progressive subfolding of the initial folds (Lawton et al., 2019). Moreover, while the development of the cerebral cortex is largely completed prior to birth, the timing of the development of the cerebellum extends significantly postnatally (Goldowitz and Hamre, 1998), and therefore is possibly more subject to environmental influences that might affect its degree and pattern of folding. Finally, the folding of the cerebellum is distinctly 2-dimensional, with folia placed mostly transversally to the midsagittal plane, in contrast to the 3-dimensional folding of the cerebral cortex. Despite so many distinctions, the cerebellar cortex, like the cerebral cortex, is a self-avoiding grey matter surface that envelops white matter and deep nuclei and has a characteristic thickness. Thus, if both cortices are self-avoiding surfaces whose expansion in development is subject to the same universal physical dictates, then the degree of folding of both cerebral and cerebellar cortices should arise from the same universal folding mechanism, regardless of the biology of how the tissue is formed – even if the spatial placement of the folds is somewhat variable, though still clade-specific. If this is the case, cerebellar morphology should be approximated by a fractal.

Fractals are geometric objects that are hierarchically composed of substructures of arbitrarily small scales (Losa 2011). These substructures are moreover self-similar, in the sense that they are scaled-up or scaled-down versions of each other, regardless of size. This self-similarity is typically quantified as a power law relation between the object’s intrinsic and extrinsic sizes, as measured by tiling it over with tiles of a fundamental, specified size. The exponent in this relation is proportional to the so-called fractal dimension of the object. Valid across mammalian cerebral cortices, the universal gyrification relation can be rewritten suggestively as 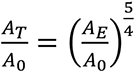, where 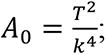; and indeed it has recently been shown (Wang et al 2022), by erasing structures smaller than a varying *A_0_*, that individual primate cortices also follow a version of this law, and thus cortices approximate fractals with fractal dimension *d_f_ = 5/2*.

Here we test the hypothesis that the cerebellar cortex folds like a fractal, following a universal scaling rule analogous to the one observed in the cerebral cortex, in a self-similar manner. We proceed by determining whether the following criteria are met. First, to be universal, folding must follow a scaling rule that applies equally across the cerebella of various species and clades. Second, there must be a fundamental unit of length, or critical length, to cerebellar folia that, once reached, is followed by formation of a new fold, in an ever-repeating pattern. Finally, scaling must follow the same scaling rule at different levels *within* an individual cerebellum. At the same time, we test the prevailing assumption that all mammalian cerebella share a common spatial pattern of distribution of their folds (and therefore lobules), as well as the alternative hypotheses that (1) there is no particular order to the placement of the folds of the cerebellum across species, and that (2) like the cerebral cortex (Welker, 1990), clade-distinctive spatial patterns of folding also occur in the cerebellum.

## MATERIALS AND METHODS

### Experimental design

We approached the investigation of whether cerebellar folding follows a universal scaling, possibly with clade-specific spatial patterns, by generating contour tracings of the total and exposed perimeters of the cerebellar grey matter as seen in the mid-sagittal section of the cerebellar vermis of different mammalian species. Because cerebellar expansion in mammalian evolution occurs most obviously at the lateral hemispheres (Kielan-Jaworowska et al., 2004), the vermis is the portion of the cerebellum most likely to be conserved in structure across species, making them most directly comparable. Additionally, the mid-sagittal vermis is readily observable in most brain hemispheres available in various brain collections, making our analysis readily feasible and easily repeatable by other groups. Our choice of quantifying folding in the midsagittal section of the cerebellum is validated by recent analyses of cerebellar folding used a similar approach (Cunha et al., 2021; Iwaniuk et al., 2006), which found that the folding index of the midsagittal section represents the folding index of the whole cerebellum (Iwaniuk et al., 2006). Importantly, the folding index calculated by those researchers uses the surface area of the Purkinje cell layer, not the pial surface of the cerebellum; here we choose to use the pial surface instead, for two reasons. First, the cerebella we use are whole, and in storage for future studies, so they cannot be stained, which precludes visualization of the Purkinje cell layer; and second, the pial surface area combined with the volume of the cerebellar cortical gray matter gives us an estimate of the average cortical thickness, which is a necessary variable for our analysis.

For the comparison of spatial patterns of cerebellar folding across species and clades, we isometrically scaled the contours so that they could be overlaid across species. Estimated variables were then subjected to mathematical analysis as described below.

### Specimens

We analyzed one specimen each of 53 species belonging to six mammalian clades: Artiodactyla (n=8), Carnivora (n=10), Marsupialia (n=5), Perissodactyla (n=3), Primata (n=8), and Rodentia (n=19; Table 1). All cerebella had the mid-sagittal surface imaged at high resolution (1200 dpi) on an HP flatbed scanner with a millimeter ruler placed on the flatbed scanner for precision and consistency.

**Table 1.**
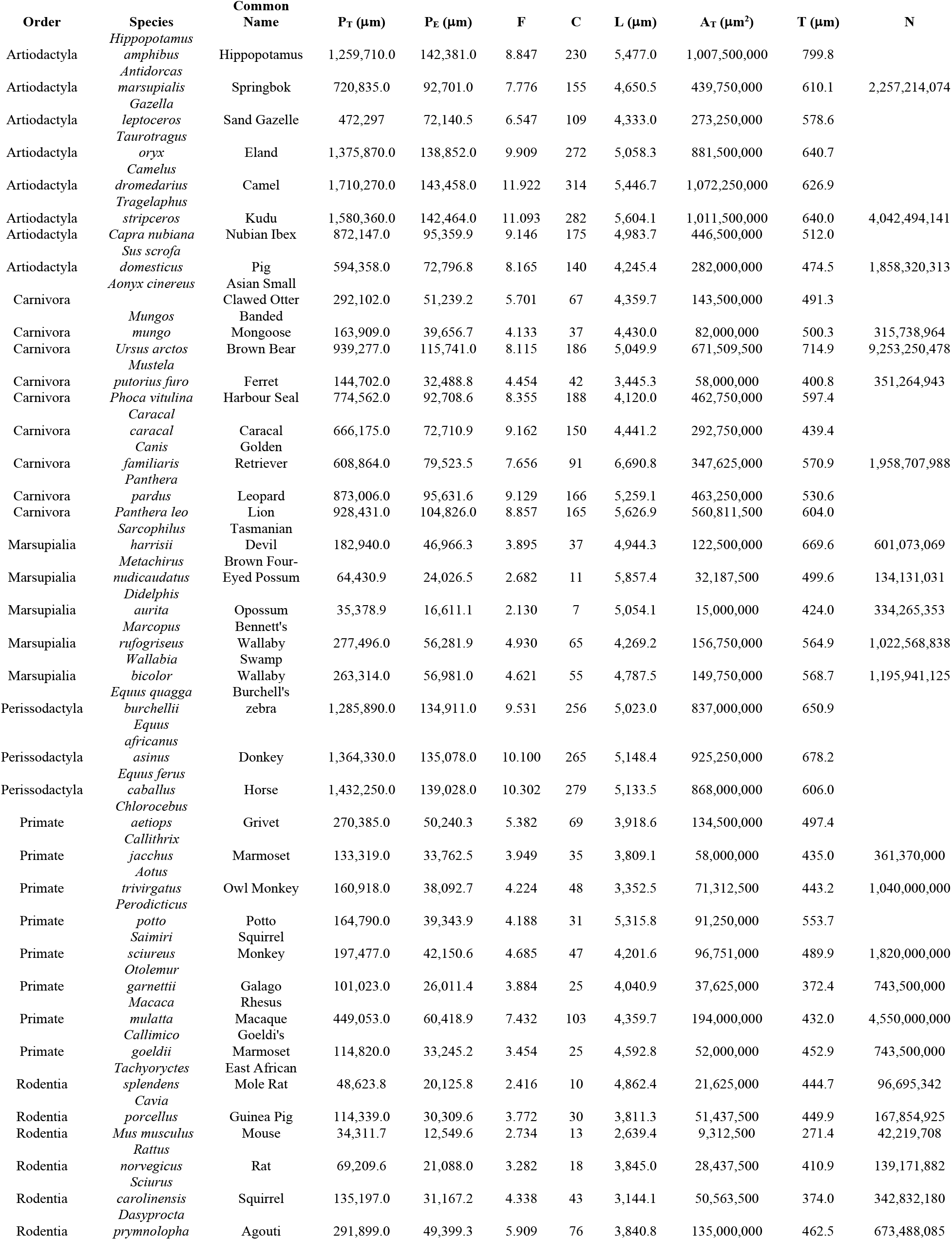
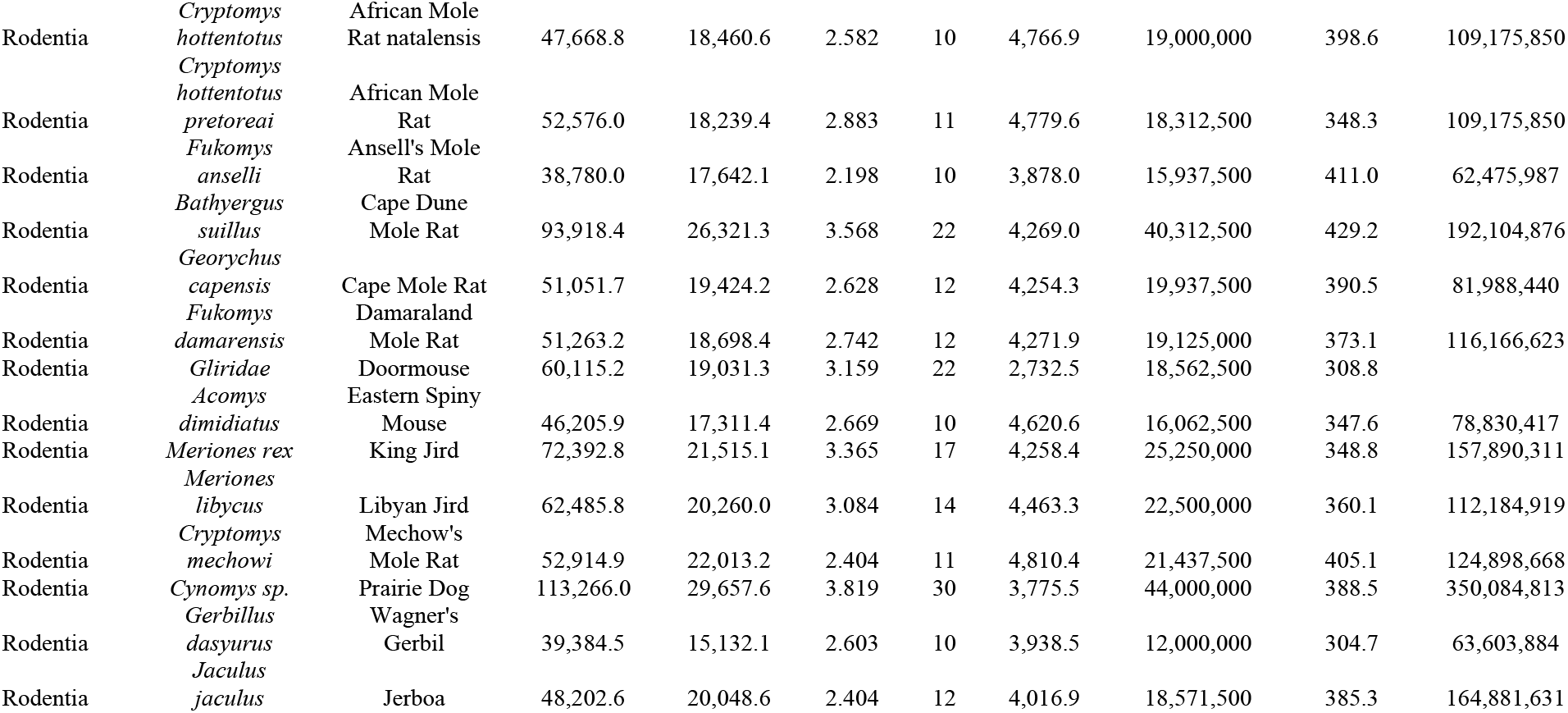
Mammalian species examined (n=53) and full dataset.

### Measurements

We used StereoInvestigator software (MicroBrightField, Williston, VT) to trace and measure all contours of the mid-sagittal cerebellar plane of each cerebellum. A virtual lens was first created to the scale of the original image for calibration. We started by tracing the total perimeter of the pial surface of the cerebellar grey matter, or P_T_, with care to capture the full contour of each folium, under enough magnification that the image of the cerebellum filled the screen of a 32” monitor (Figure 1, P_T_). Next, a new contour traced the outer (exposed) surface of each cerebellum, from the anterior-most to the posterior-most edge of the cerebellar cortex, as if saran-wrapping the structure, to provide a measurement of the exposed perimeter (P_E_) of the midsagittal cerebellar plane (Figure 1, P_E_). The degree of folding in this two-dimensional plane, or folding index F, was calculated for each cerebellum as the ratio between the total perimeter and the exposed perimeter of the cerebellar grey matter on the midsagittal plane (F = P_T_ / P_E_).

**Figure 1.**
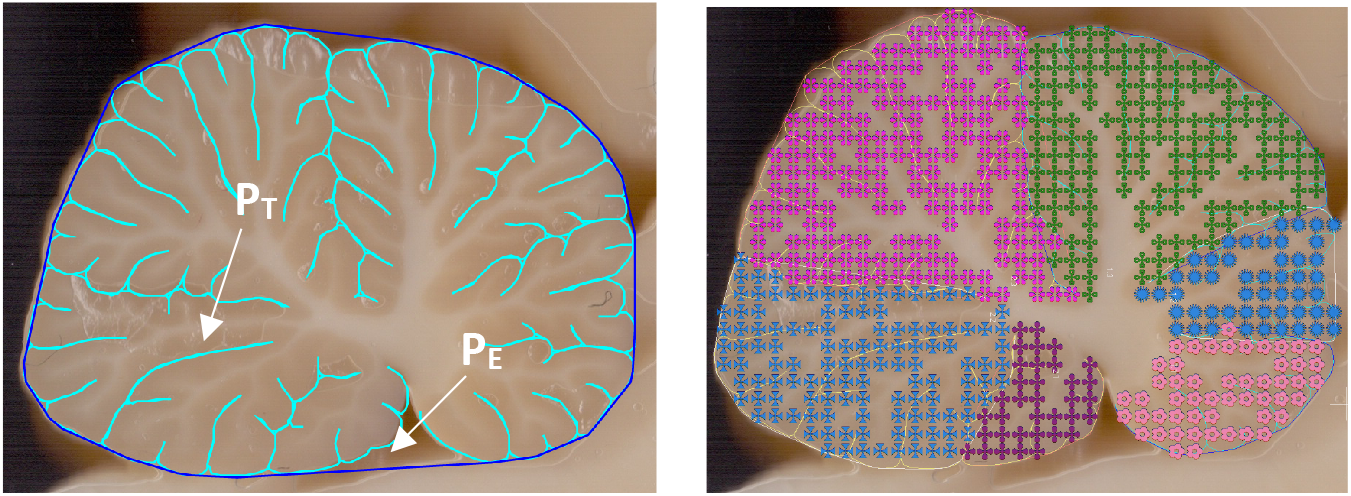
Contour tracing of total and exposed perimeters, and Cavalieri estimation of sagittal surface area for estimating grey matter thickness. **Left,** Contours of total grey matter perimeter P^T^ (light blue) and exposed grey matter perimeter P_E_ (dark blue), from which folding index F is calculated as P_T_/P_E_. **Right,** Cavalieri probe used to obtain sagittal surface area AT for estimation of average grey matter thickness T = A_T_/P_T_ for each subdivision and for the whole sagittal surface. All measurements were made using StereoInvestigator software.

To estimate the average thickness of each cerebellar cortex, we utilized the Cavalieri probe within StereoInvestigator to obtain measurements of the area of grey matter on the midsagittal plane (A_T_), which when divided by the total perimeter provides a measurement of the average grey matter thickness (T = A_T_ / P_T_) at that plane. Next, we determined the number of folia in the midsagittal plane (C) by counting how many foliar crowns were contained in P_T_, which allowed us to determine the average length L of the cerebellar folia in each species as the ratio L = P_T_/C (Figure 1, right). While we define a “foliar crown” as the unit of cortical surface farthest from the white matter in any segment of cerebellar cortex between two sulci (that is, a gyrus) in the recurring folding of this cortex. Note that this procedure does not require or involve measuring the length of individual folia in order to determine the average length of cerebellar folia, L.

### Mathematical model

Perimeter measurements of the midsagittal surface of the cerebellum were used rather than estimates of the full surface area of the cerebellar cortex because the latter would be too time-consuming, and actually unnecessary given the 2-dimensional nature of cerebellar folding, which makes the FI of the midsagittal plane (P_T_/P_E_) a good proxy for the FI of the entire cerebellar surface (A_T_/A_E_) (Iwaniuk et al., 2006). Importantly, the model of Mota and Herculano-Houzel (2015) can be adapted to the P_T_ x P_E_ relationship as follows, with the only caveat being that the predicted exponent for the universal function becomes 1.5 (rather than 1.25), similar to what was found to apply for coronal sections of the cerebral cortex (Mota & Herculano-Houzel, 2015). We thus predict that for the cerebellum, as for coronal sections of the cerebral cortex, P_T_.T^1/2^ = k.P_E_^1.5^.

Our previous description of a simple mechanism for cerebral cortex folding was based on the self-avoiding nature of cortical surfaces, on one hand, and on the elongation/contraction response of white matter axons under longitudinal stresses (effectively viscoelastic over appropriate time scales), on the other. The resulting model (Mota and Herculano-Houzel, 2015) implies that the shape of the adult cerebral cortex is an equilibrium configuration that minimizes an effective free energy: A combination of a low-enough effective energy associated with axonal length, and a high-enough configurational entropy associated with a crumpled surface, where the exchange rate between the two is defined by the effective temperature, τ. In that model, we used a so-called Flory approximation and obtained that a cortex intrinsically characterized by a total area A_T_ and thickness T, and in a configuration with exposed area A_E_, will have energy 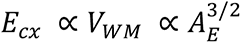 and entropy 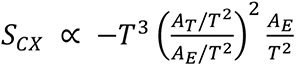.

Thus, minimizing free energy *F*_*CX*_ = *E*_*CX*_ − τ *S*_*CX*_ with respect to A_E_, we get 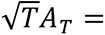 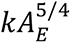, where 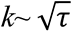 is a universal dimensionless parameter not specified by the model. This latter relation is in close agreement with empirical results for cortices across both species and individuals (Mota and Herculano-Houzel, 2015, Wang *et al*, 2016, Wang *et al*, 2018), and was shown to also be valid across length scales for individual primate cortices (Wang *et al*, 2022), indicating cortical shape is the self-similar hierarchical aggregation of structures over a range of length scales (and thus approximately fractal). In this context, k can be reinterpreted as defining the low end of this range: the near-invariant typical area of the smallest sulci and gyri present in cortices, that is, α_0_, defined as a multiple of cortical thickness squared. The value of α_0_ can be obtained directly from the universal relation by calculating the area *A_0_* for which the transition between lissencephaly and gyrencephaly occurs, i.e., *A_0_* = *A_T_* = *A_E_*, then dividing *A_0_* by *T^2^*, in which case we obtain that α_0_ ≅ 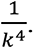

The above description characterizes the outer boundaries of white and gray matter in the cerebral cortex as nested two-dimensional self-avoiding surfaces that fold in three-dimensional space. In principle, the cerebellar cortex could be described in the same manner. However, due its peculiar two-dimensional folding pattern, discussed above, we posit that a medial sagittal section of a cerebellum will capture most of the relevant information concerning how it folds. Thus, we modelled cerebellar folding as that of nested self-avoiding contours that fold in two-dimensional space.

Using the same framework as before, a cerebellar slice with total perimeter P_T_ and thickness T, and in a stable equilibrium configuration with exposed perimeter P_E_, will have energy 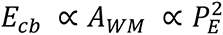 and entropy 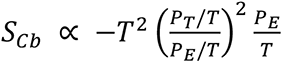. Thus, minimizing free energy *E*_*CX*_ = *E*_*CX*_ − τ *S*_*CX*_ with respect to P_E_, we get the theoretical scaling relation 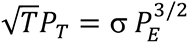, where σ is a universal dimensionless parameter not specified by the model. This is the scaling relation we seek to test empirically.

To test for self-similarity, we determine if the same scaling rule applies at each hierarchical level within the cerebella (from the whole contour to individual lobules). We thus perform the same scaling analysis and measure all variables for the mid-sagittal cerebellar surface as a whole (level 0, which encompasses the entirety of the white matter within its contour); the two halves of the midsagittal plane divided by the main (deepest) cerebellar fissure (level 1); as well as increasingly smaller subdivisions of each, down to lobules (level 2 and beyond; Figure 2).

**Figure 2.**
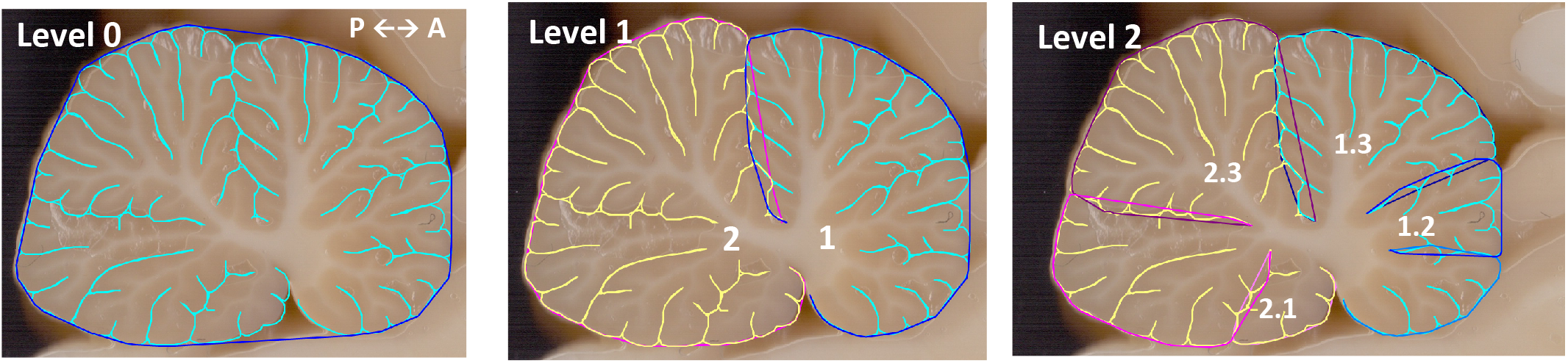
Hierarchical division levels of pial surface contours for data analysis, shown from Level 0 (whole contour) to Level 2. Species shown: *Aonyx cinereus* (Asian small clawed otter). Notice that the contours of progressive subdivisions include less and less of the subcortical white matter.

Finally, the hypothesized theoretical scaling relation would be strongly suggestive of fractal scaling with an invariant fundamental scale *p*_0_ for the typical length of the smallest structures, with 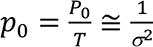 (as a multiple of thickness), where *P_0_* is an estimate for average foliar length. Empirically, we compute the distribution of average cerebellar foliar lengths (calculated as above) for each species, and compare each species’ average foliar length to the expected scale derived from the analysis outlined above to look for evidence of an invariant fundamental scale across folia of all species.

### Scaling analysis

In addition to the morphological variables P_T_, P_E_, A, T, C and L, we estimated numbers of neurons in the cerebellum, N_CB_, by using previously obtained equations relating N_CB_ to brain mass, which excludes perissodactyls (Table 2). We then analyzed the scaling relationships across all variables, and tested for the power law 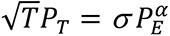 by using least squares regression fitting to power functions using JMP 14 (SAS, Cary, NC), where α=3/2 is the exponent predicted by our folding model.

**Table 2.** Equations used to estimate number of cerebellar neurons from brain mass, by clade.

| Clade | Predictive Equation |
| --- | --- |
| Artiodactyla | $N_{cb} = e^{18.207 \pm 0.683} M_{br}^{0.714 \pm 0.130} r^2=0.909 p<0.0001$ |
| Carnivora | $N_{cb} = e^{17.907 \pm 0.446} M_{br}^{0.828 \pm 0.108} r^2=0.908 p<0.0001$ |
| Marsupialia | $N_{cb} = e^{18.326 \pm 0.103} M_{br}^{0.747 \pm 0.038} r^2=0.982 p<0.0001$ |
| Primata | $N_{cb} = e^{18.297 \pm 0.190} M_{br}^{0.876 \pm 0.048} r^2=0.974 p<0.0001$ |
| Rodentia | $N_{cb} = e^{18.272 \pm 0.104} M_{br}^{0.672 \pm 0.051} r^2=0.955 p<0.0001$ |

### Spatial patterning of folding

The spatial arrangement of the main cerebellar lobes and folia were qualitatively compared across the 53 species by aligning the contours traced from the midsagittal plane for all species. Each traced contour was copied from StereoInvestigator over to Adobe Illustrator (Adobe, San Jose, CA), and all contours were scaled isometrically to match an arbitrary length, to make them comparable. The contours were then superimposed in series of within-clade and across-clade comparisons to determine the degree of overlap of the main lobes. For each of these overlay analyses, the contours were aligned following a set of predefined criteria. First, all were placed in the same anterior-posterior, ventral-dorsal orientation. Next, they were overlaid upon one another by matching the anterior and posterior edges of the cerebella, then rotating one image to align its main, deepest fissure (the fissure which divides the cerebellum into two relative halves, whether or not that is called the primary fissure and the lobes it creates, the anterior and posterior cerebellar lobes), to the reference image. The final placement of each cerebellum sought to maximize overlap across sulci and, where possible, lobules and folia. To facilitate detection of coincidence of the main lobes across species, outline traces were also taken that followed only the main sulci for each cerebellum, ignoring lobules and folia, and similarly overlaid both within-clade and across-clade.

## RESULTS

Contours of the midsagittal plane of the 53 cerebella arranged in order of increasing total perimeter (P_T_) of the cerebellar cortical surface demonstrate the wide diversity encompassed within the study sample (Figure 3) – from the mouse cerebellum, with a P_T_ of only 34 mm, to the large and highly folded cerebellum of the camel with a P_T_ of nearly 2000 mm, a range of 59-fold. Yet, despite the diversity in total cerebellar mid-sagittal perimeter represented in the sample, the individual folia of each cerebellum appear to be of comparable lengths (quantified below).

**Figure 3.**
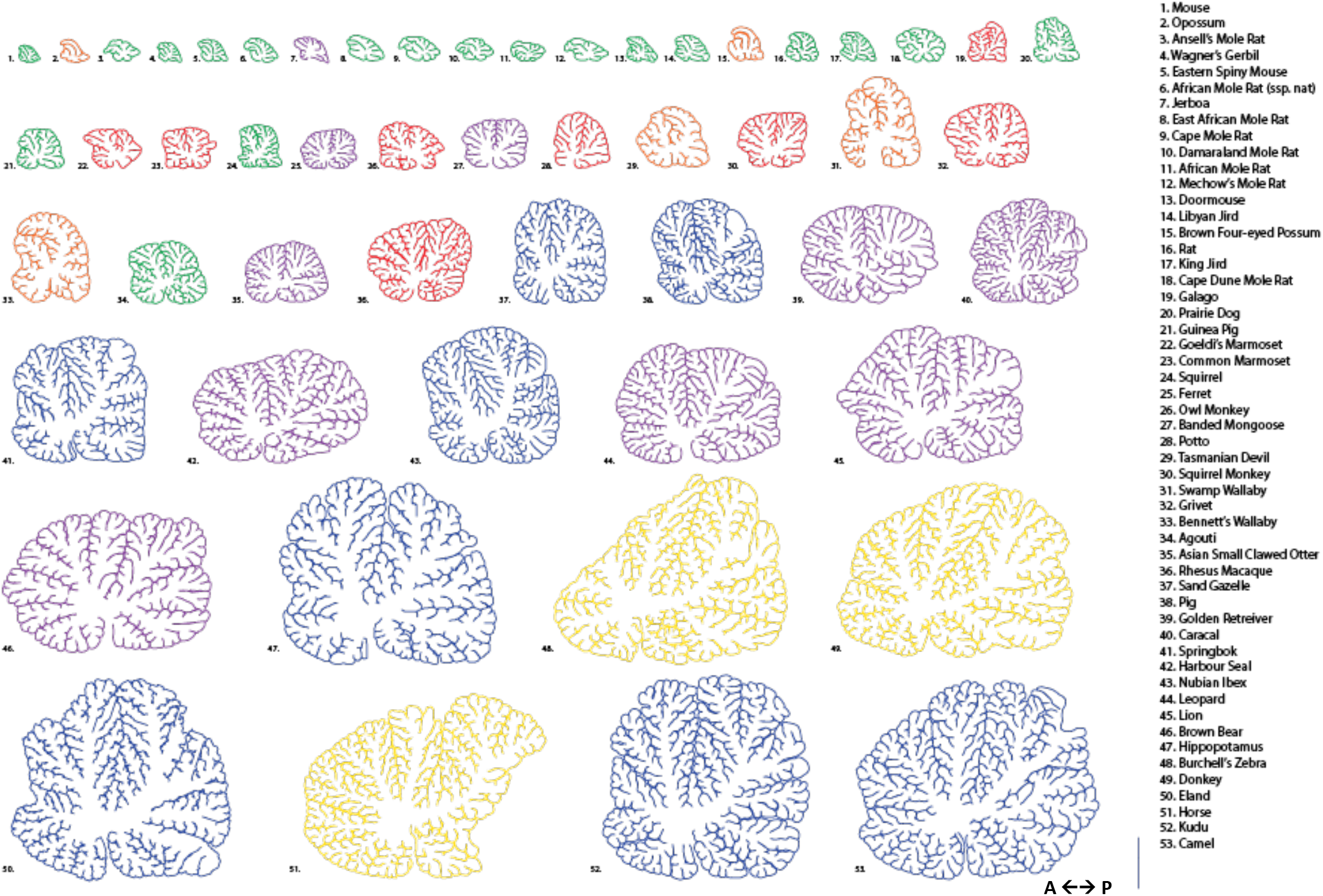
Array of 53 species mid-sagittal pial contours, in order of increasing P_T_ and colored according to clade. Contours are colored as follows: Artiodactyla in blue, Carnivora in purple, Marsupialia in orange, Perissodactyla in yellow, Primata in red, and Rodentia in green. Scale bar, 1 cm.

It is also clear from the all-species array depicted in Figure 3 that there is large variability in the morphological appearance of the cerebellar contours and the spatial arrangement of their lobes and folia across species. However, color-coding for the different clades reveals patterns in the spatial arrangement of cerebellar lobules in that species of the same clade have similar overall distribution of cerebellar folds (Figure 3).

These clade-specific spatial patterns become more visibly apparent in a comparative arrangement of cerebella of species organized by increasing size (i.e. total perimeter length) within each clade, as illustrated in Figure 4. For example, the cerebella of the artiodactyls appear relatively taller than others, with distinct branching fingers of white matter that penetrate the highly folded grey matter (Figure 4A). The carnivoran cerebella, in contrast, seem to expand more horizontally, wider than they are tall (Figure 4B), while the primate cerebella are more evenly square (Figure 4D), and marsupial cerebella are decidedly asymmetrical, possessing a distinctive “hook” that faces anteriorly (Figure 4C). In fact, the patterns are so consistent among species within each clade that it appears visually as though they were scaled-up or scaled-down versions of the same cerebellum, following the same folding pattern as it expanded in development.

**Figure 4.**
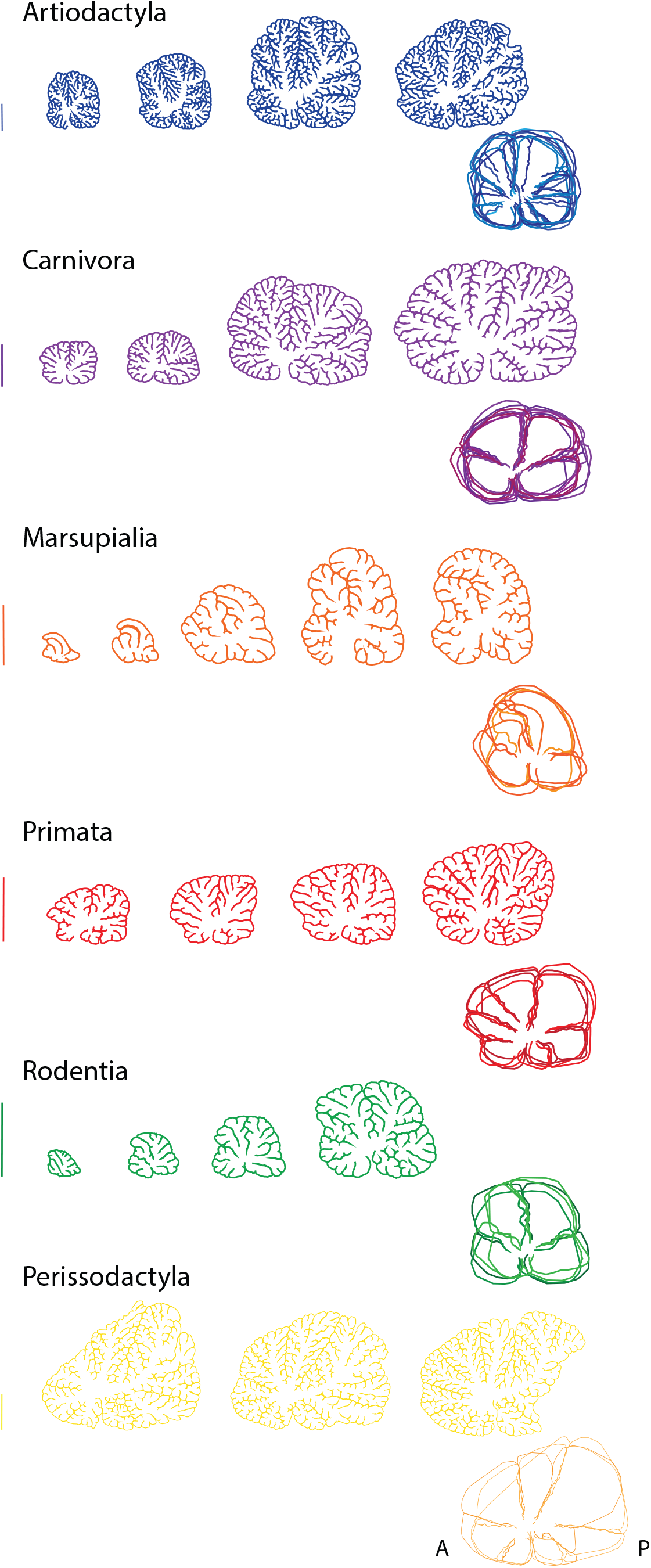
Array of select contours for each clade, arranged in order of increasing P_T_. Artiodactyla: *G. leptoceros, C. nubiana, T. stripsceros, C. dromedarius*. **Carnivora:** *M. mungo, A. cinereus, P. pardus, U. arctos*. **Marsupialia:** *D. aurita, M. nudicaudatus, S. harrisii, W. bicolor, M. rufogriseus*. **Primata:** *A. trivirgatus, S. sciureus, C. aetiops, M. mulatta*. **Rodentia:** *Mus musculus, Rattus norvegicus, C. porcellus, D. Prymnolopha*. **Perissodactyla:** *E. quagga, E.asinus, E. caballus*. Scale bar, 1 cm.

With the contours arranged by size per clade in Figure 4, one can better appreciate that the individual folia all appear to be of comparable length across species, whatever the total perimeter of the cerebella or its degree of folding, which we explore quantitatively below. This observation is in line with the expectation that if cerebellar folding is fractal, then once the developing folia grow in length to a certain multiple of the cortical thickness, they fold in on themselves forming two new folia of approximately equal perimeter length. Further, overlaying the outer contours of the cerebella of the select species as depicted in Figure 4 reveals alignment of the contours within each clade, which indicates a constant spatial pattern of folding, regardless of cerebellar size.

### Clade-dependent spatial arrangement of main cerebellar lobes

To investigate the conservation of the spatial distribution of the main folds of the cerebellum, we overlaid contours traced exclusively along the main sulci in the cerebellum of each species, isometrically scaled to similar linear dimensions. Figure 5 demonstrates the clear and consistent overlap in the shape of the main cerebellar halves and quarters across all species examined within each clade. Figure 6 shows the overlain full pial contours of all species within each clade, in which clear overlap of not only the main halves and quarters but also the folia becomes evident. For clarity, the individual contours in the overlays are shown separately, once isometrically scaled for comparison, in Figure 7.

**Figure 5.**
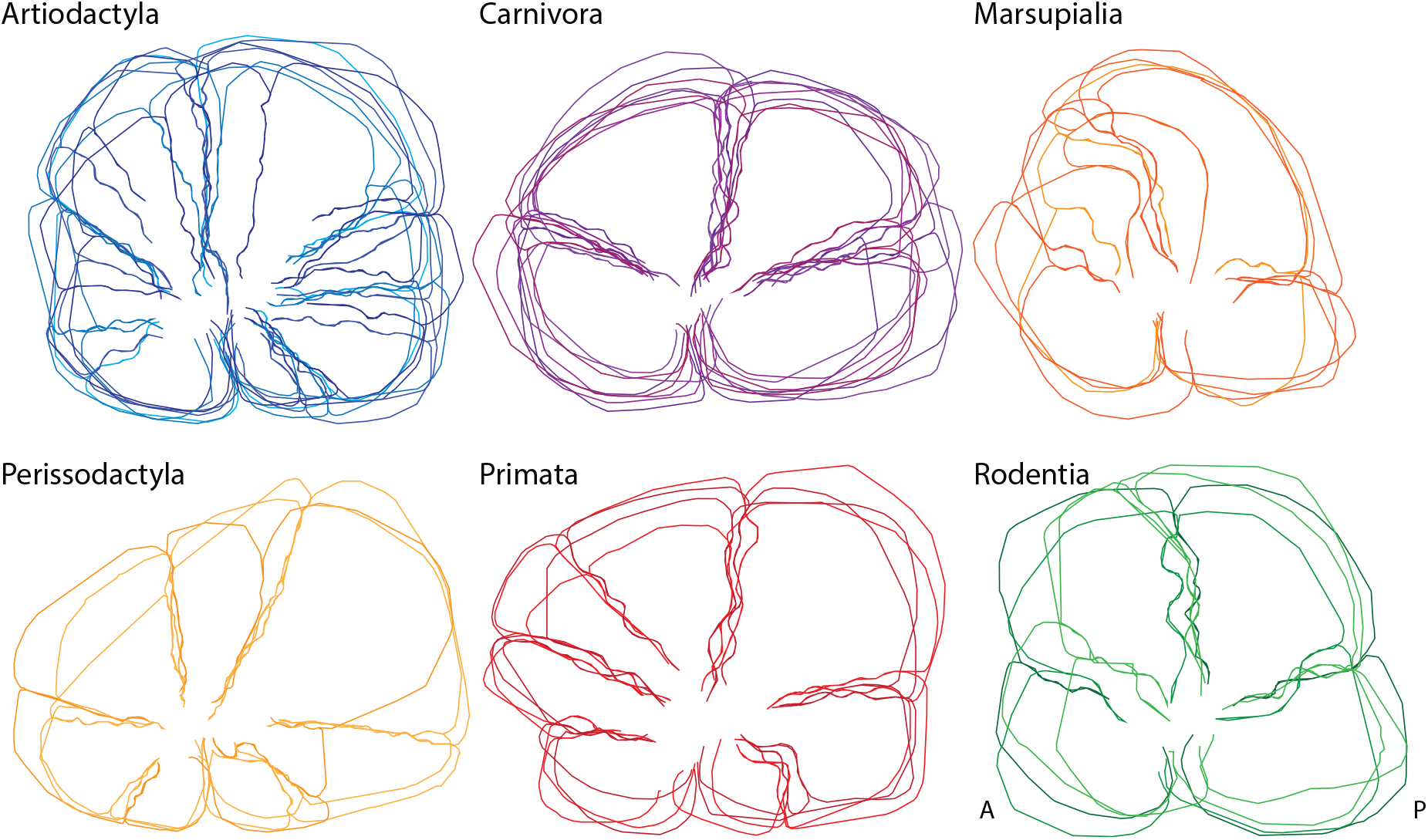
Overlay of main sulci contours of the mid-sagittal surface of the cerebellum demonstrates nearly identical spatial arrangement of main lobes within clades. A, anterior; P, posterior.

**Figure 6.**
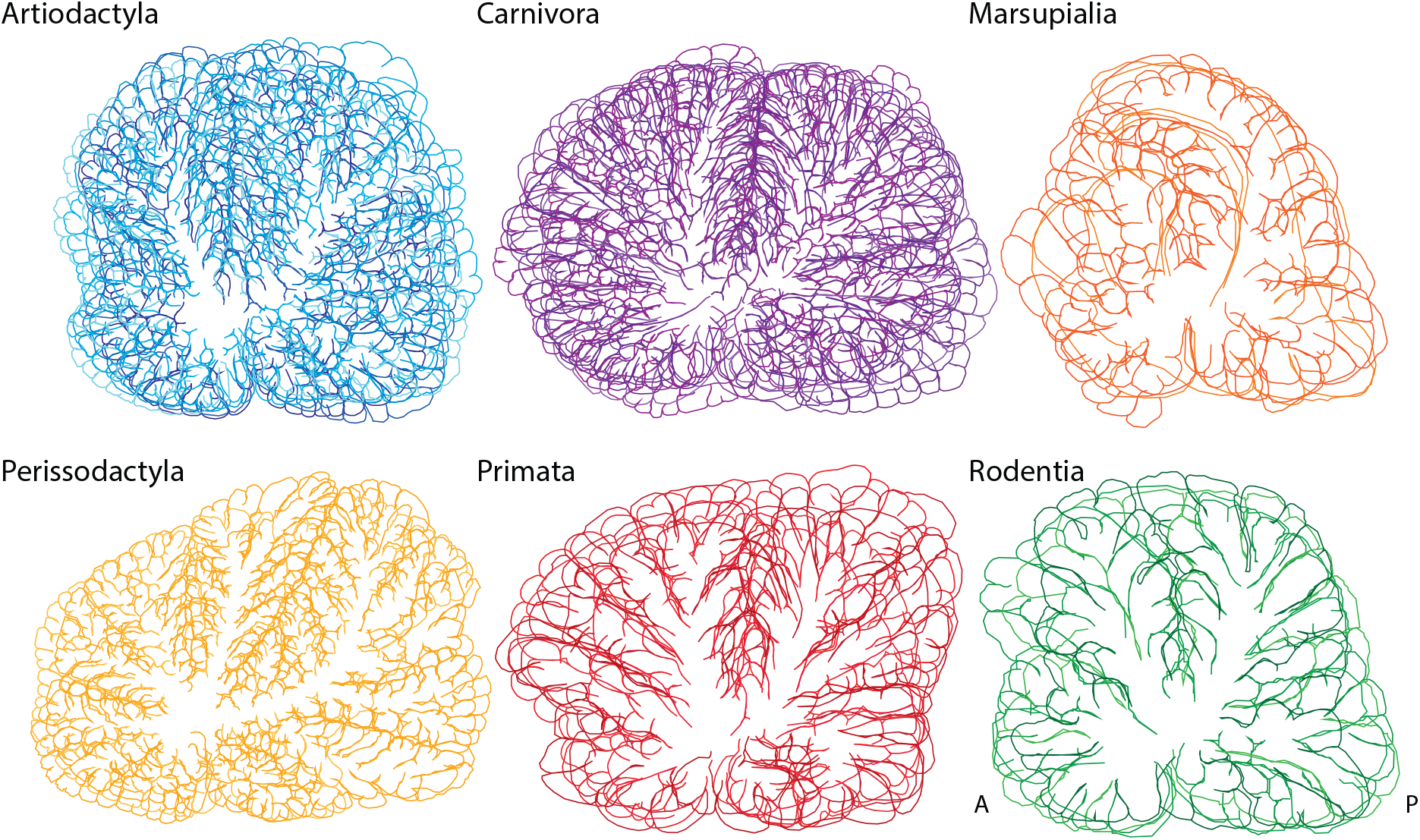
Overlay of full contours of the mid-sagittal surface of the cerebellum demonstrates consistent overlap of the location of main lobes and folia across species within each clade. A, anterior; P, posterior.

**Figure 7.**
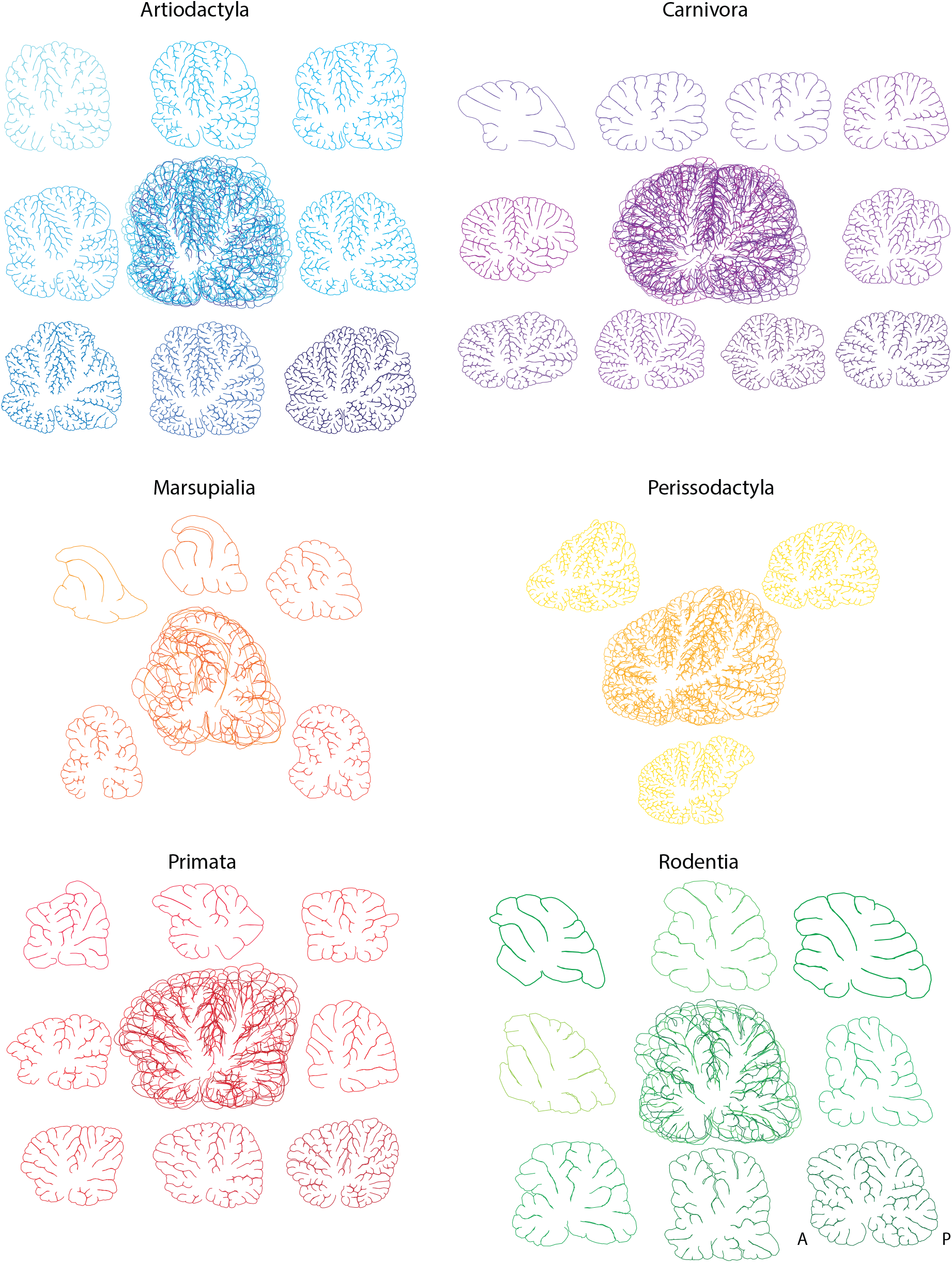
Side-by-side display of individual full contours of the mid-sagittal surface of the cerebellum with overlays for each clade demonstrates consistent overlap of the location of main lobes and folia across species within each clade. A, anterior; P, posterior.

In contrast, it is difficult to obtain a similar overlay of the full pial contours for cerebella of species belonging to different clades, demonstrating an inconsistency in the spatial arrangement patterns across clades (Figure 8, left). Interestingly, and again consistent with hierarchical, fractal folding of the cerebellar surface, an all-species overlay of just the main sulci contours does show some consistency, as seen in Figure 8 (right). Although the overlap of the main, central fissure is a given, since it was used as the point of alignment for all contours, it is remarkable that alignment of the main fissure across all species also allows good alignment of the anterior and posterior borders of the perimeters. Thus, the excellent alignment of the main fissure together with the anterior and posterior extremes of the cerebellum across all species indicates that the main fissure is consistently placed dividing the midsagittal plane of the cerebellum into halves. Next, there is relatively consistent overlap of the two somewhat horizontally oriented sulci that further divide each cerebellar half into quarters, but after this second division, the remaining fissures become increasingly inconsistent in placement. This arrangement is indicative of hierarchical folding, which will be discussed subsequently. Overall, these results contradict the common expectation that all mammalian cerebella share the same conformational plan, including subdivisions into similar and corresponding lobules. Rather, it is evident that each mammalian clade has a distinctly unique spatial arrangement of its cerebellar folding pattern.

**Figure 8.**
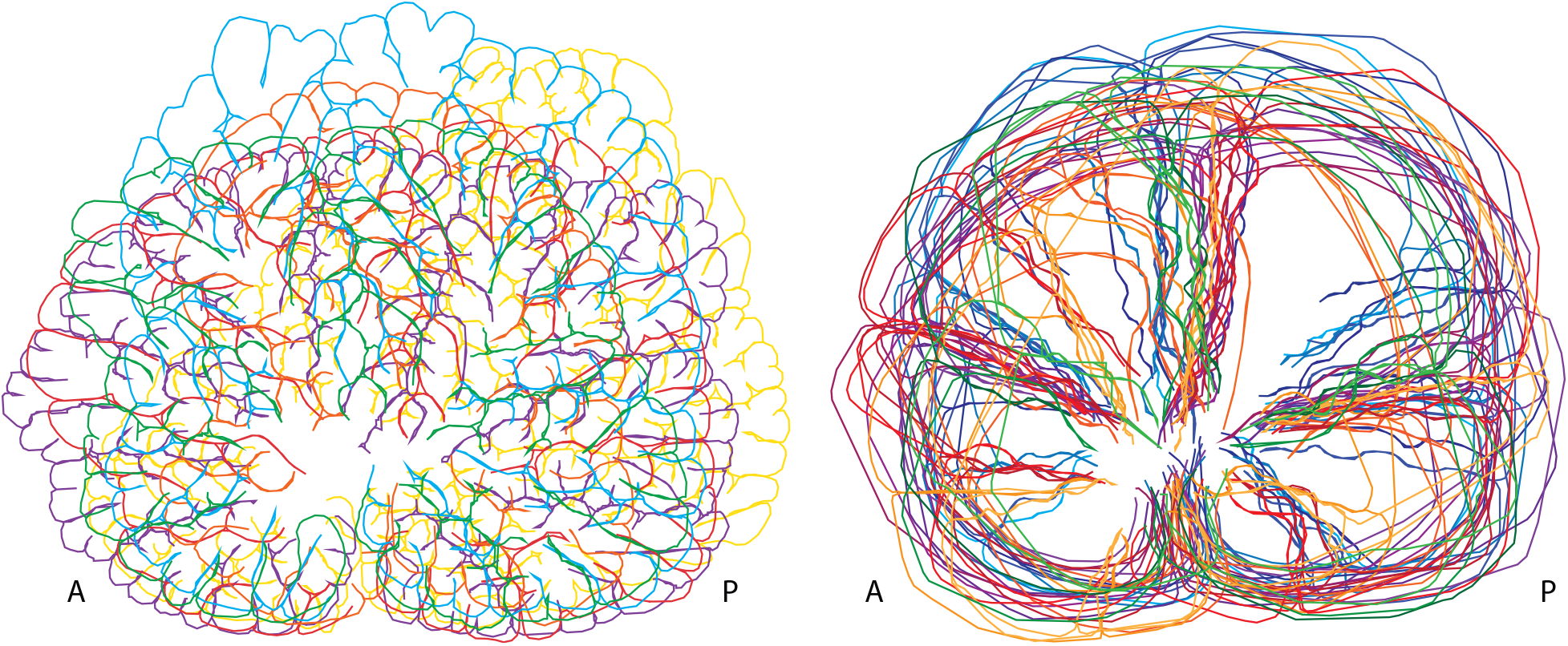
Overlay of the mid-sagittal pial contour of species belonging to different mammalian clades demonstrates inconsistent overlap of the folia (left) but somewhat consistent overlap of the main sulci (right). Selected species from each mammalian clade: *G. leptoceros* (artiodactyl, blue)*, U. arctos* (carnivoran, purple)*, M. rufogriseus* (marsupial, orange)*, E. quagga* (perissodactyl, yellow)*, S. sciureus* (primate, red)*, and D. prymnolopha* (rodent, green).

### Hierarchical folding of the cerebellum

Hierarchical folding is expected of structures that fold as a fractal, but it is not a sufficient condition for fractality. If the cerebellar cortex folds hierarchically, then the first fold should form at an approximately central location that should also be the deepest, with subsequent folds forming sequentially at the approximate halfway point of the previously formed fold. To determine if this is the case, we identified the cerebellar sulci in the midsagittal plane of each species, labeled them hierarchically by degree of branching order, and examined their relative depth. As illustrated in Figure 9, we find that the main sulcus which divides the cerebellar pial contour approximately in half (labeled 1) is always the deepest, while the next two deepest are the sulci that divide the perimeter of each previously formed half in half again (labeled 2). The next four deepest sulci are those that divide each quarter into eighths (labeled 3), with the number of divisions in half depending on the size and degree of folding of the individual cerebellum, as illustrated in Figure 10.

**Figure 9.**
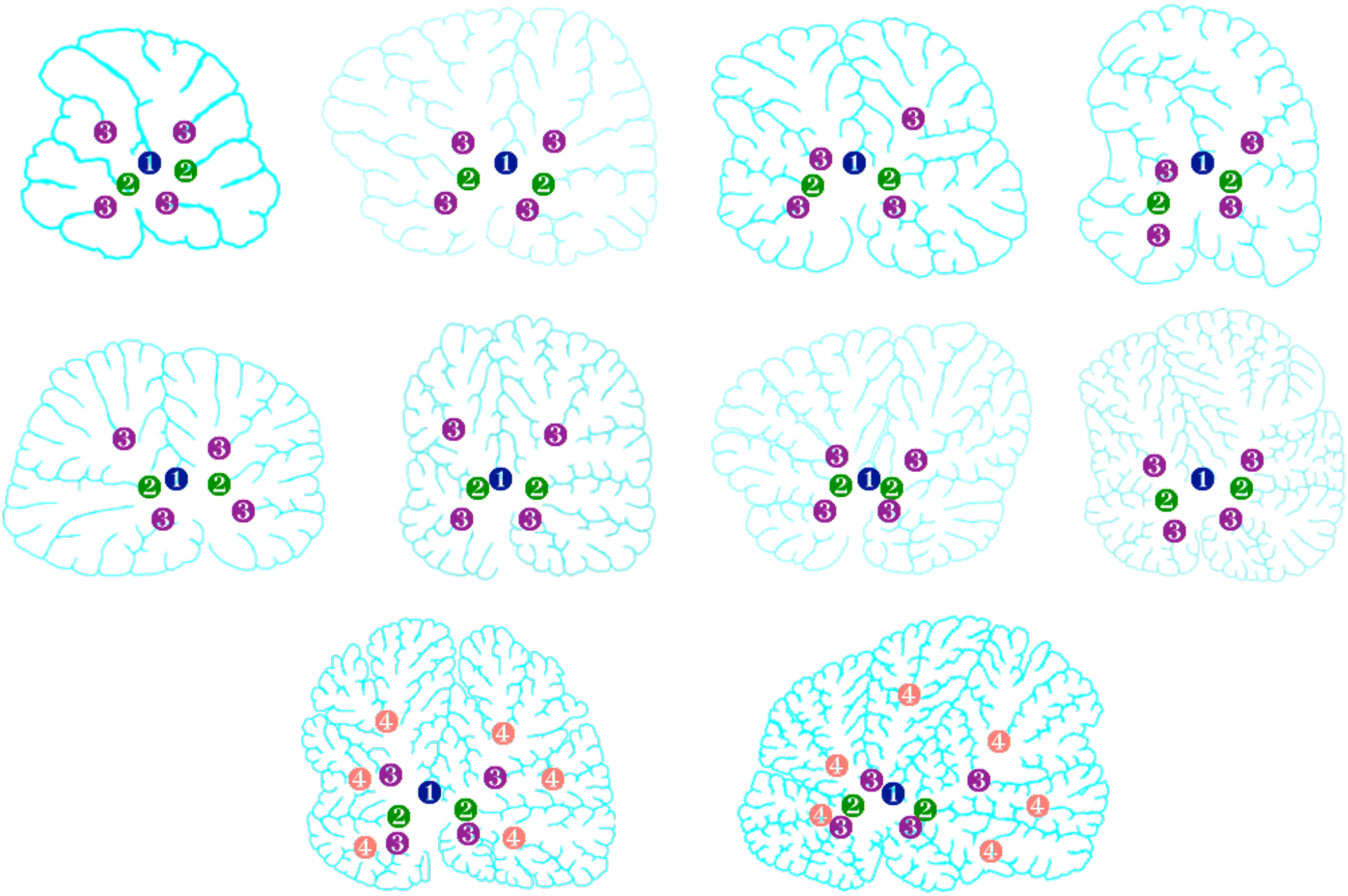
Hierarchical folding of cerebella. Across species and clades, a pattern emerges with the deepest sulci dividing the perimeter of the cerebella approximately in half, with subsequent sulci of decreasing depth each dividing the previous perimeter in half again. Species: *C. porcellus, C. aetiops, D. prymnolopha, M. rufogriseus, A. cinerus, G. leptoceros, M. mulatta, C. nubiana, H. amphibus, E. asinus*.

**Figure 10.**
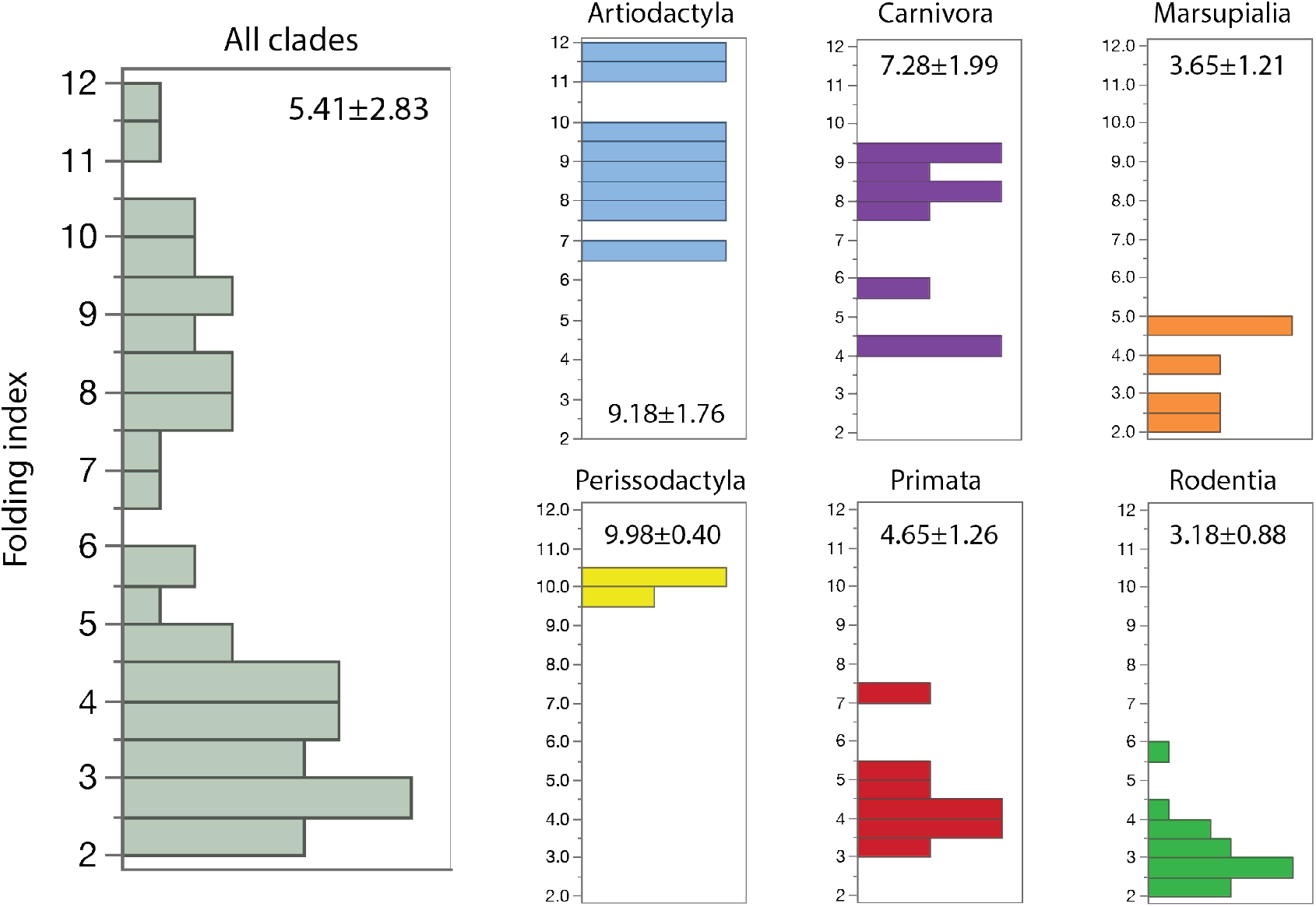
Distribution of cerebellar folding indices across mammalian species in the dataset. Values for all 53 species in the dataset, ranging from 2.13 to 11.92, are shown in the histogram on the left, and for each clade separately to the right, color-coded as in the other Figures. Average folding index ± standard deviation for each distribution is shown in the respective histograms.

These results support a hierarchical pattern of cerebellar folding that is qualitatively consistent with fractal folding. We next performed a quantitative analysis of cerebellar folding to determine whether variations in the degree of folding across species also follow a fractal pattern.

### Diversity in degree of folding of the cerebellar surface

The folding index of the cerebellar cortex ranges from 2.13 to 11.92 across the 53 mammalian species, with an overall mean of 5.41 ± 2.83 (Figure 10). The larger the perimeter of the mid-sagittal cerebellar section, the more folded the cerebellar surface (Figure 11A; r^2^=0.978, p<0.0001); artiodactyl, perissodactyl and carnivoran cerebella, the largest in the sample, are clearly the most folded.

**Figure 11.**
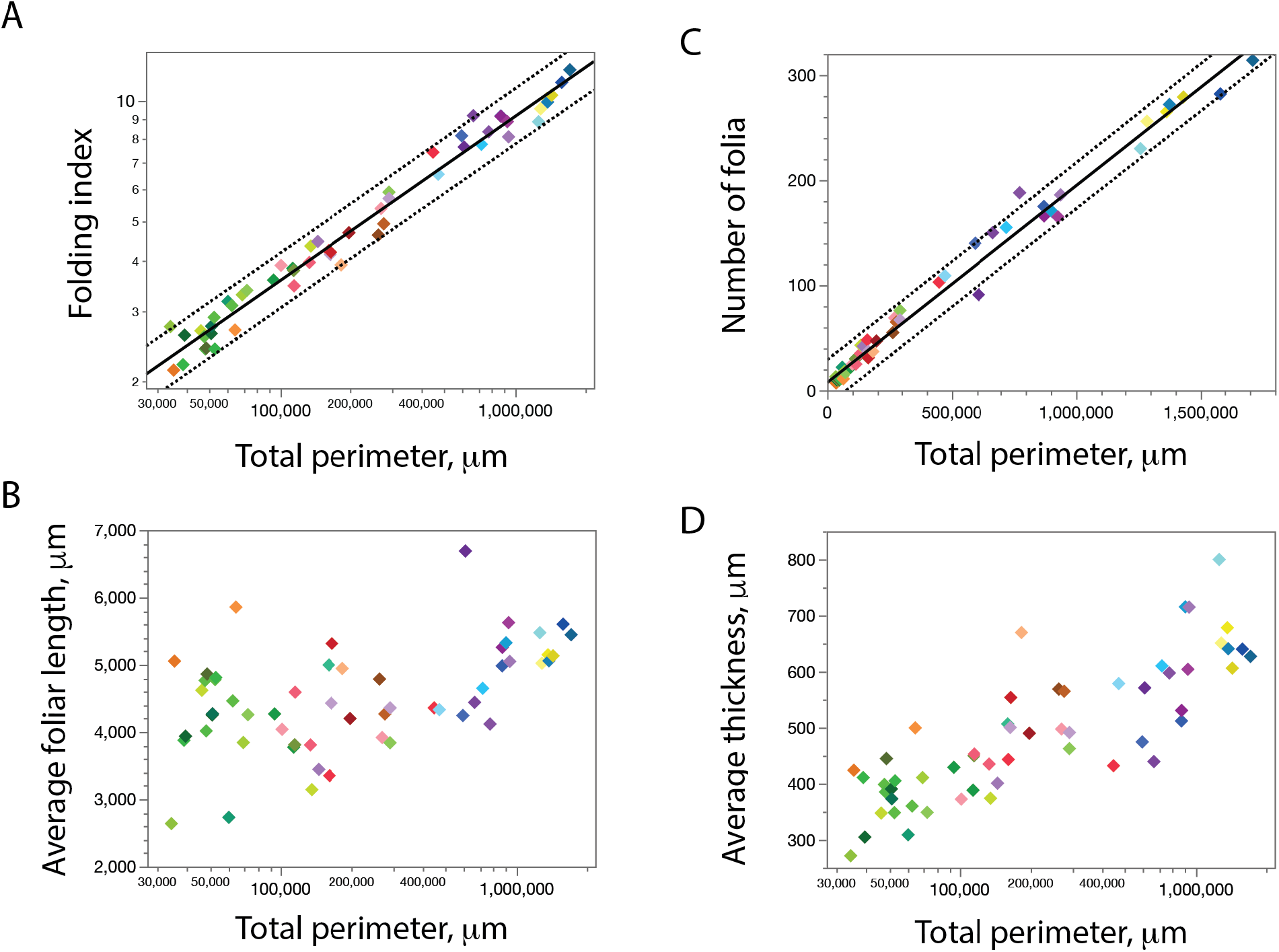
Scaling of cerebellar folding index, average foliar length, number of folia, and cortical thickness with perimeter of the pial surface of the mid-sagittal section across mammalian species in the dataset. Each point represents one of the 53 species in the dataset, color-coded in different shades according to the other Figures. Spearman correlation coefficients: 0.978 (A), 0.493 (B), 0.983 (C), 0.854 (D; all p<0.0001). Although the correlation between average cerebellar cortical thickness and total cerebellar perimeter is significant across all 53 species, it does not reach significance within each clade (artiodactyls, p=0.0876; carnivorans, p=0.0072; marsupials, p=0.2848; perissodactyls, p=0.6667; primates, p=0.3851; rodents, p=0.0733). The power function plotted in A has an exponent of 0.409±0.009 p<0.0001) and r^2^ of 0.978. Although this is consistent with universal scaling of cerebellar folding as the cerebellar surface expands, note that the midsagittal section of marsupial cerebella (orange shades) is consistently less folded than that of rodent cerebella (green shades) with similar perimeter. The linear function plotted in C has an r^2^ of 0.986 (p<0.0001), consistent with scaling of the cerebellar surface without significant changes in average length of individual folia.

### Universal scaling of degree of cerebellar folding

We find that the relationship between total and exposed surface perimeter of the cerebellar cortex, taking cortical thickness into account as predicted by our model, is a power law of form P_T_ T^1/2^ = σ P_E_^1.817±0.016^ that accounts for 99.6% of the variation in the data across all species (Figure 12A), which is more than the 97.8% accounted for without taking cerebellar cortical thickness into consideration (Figure 11A). On the other hand, the degree of folding of the midsagittal cerebellar plane is not universally predicted by the number of neurons in the cerebellum estimated from cerebellar mass (Figure 12B). While a significant correlation is evident (r^2^=0.807, p<0.0001), we find that artiodactyl cerebella are consistently more folded than primate and marsupial cerebella of similar numbers of neurons, which is compatible with the thinner cerebella of the former (see below). Similarly, for equally folded cerebella, there are systematically more neurons in those of primates and marsupials compared to rodents (Figure 12B). We thus propose that while cerebella with more neurons tend to be more folded, this correlation is not causal, but is actually a consequence of clade-specific correlations between numbers of cerebellar neurons and cerebellar perimeter.

**Figure 12.**
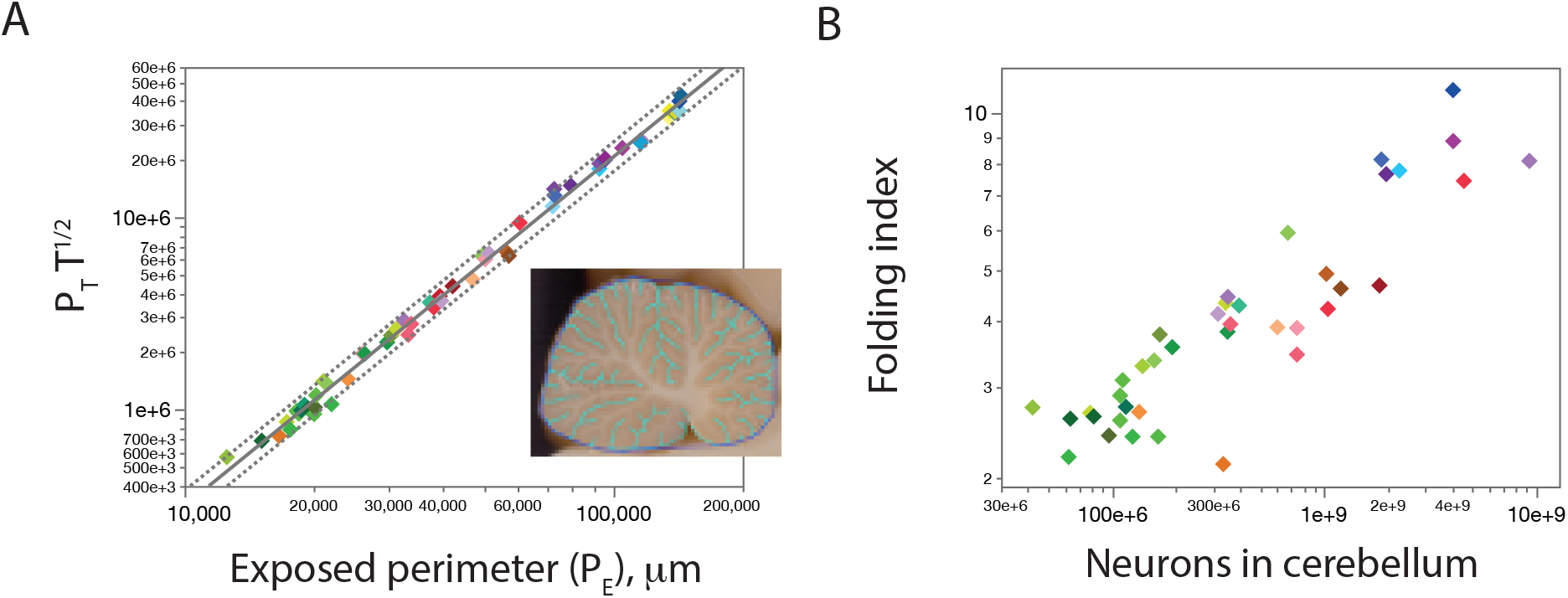
Universal scaling of cerebellar folding according to our model from first physical principles, but not with numbers of cerebellar neurons. Values for all 53 species in the dataset are shown in the graphs, color-coded as in the other Figures (shades of blue, artiodactyls; purple, carnivorans; orange, marsupials; yellow, perissodactyls; red, primates; green, rodents). The power function plotted in A has an exponent of 1.817±0.016 (r^2^=0.996, p<0.0001). For B, the power function (not plotted) has an r^2^ of 0.807 (p<0.0001), but clearly distinct relationships apply to primates and marsupials compared to other clades.

While the relationship between P_T_ and P_E_ of the whole midsagittal plane is very well explained by the equation above that takes cerebellar cortical thickness into consideration, the exponent of this equation notably does not match the value of 1.5 expected for a 2-dimensional folding pattern according to our gyrification model. Although this discrepancy may simply be due to the mid-sagittal section being an imperfect proxy for the full cerebellar surfaces, the precision of the empirical power law suggests that this reflects some universal mechanism for cerebellar gyrification.

Geometrically, could this power law be evidence that the cerebellar mid-sagittal sections are fractal with *d_f_ = 1.817*? To answer this question, and to further explore the cerebellar shape, we now test it for self-similarity. We apply our analysis to progressively higher hierarchical levels of folding, which progressively exclude the core of the midsagittal white matter. Figure 13 shows that the value of the exponent of the relationship between P_T_ T^1/2^ and P_E_ decreases progressively as analysis is moved to higher levels. At the highest level of division (level 3), where white matter contribution is minimal, the exponent is 1.501 ± 0.029, identical to the predicted value of 1.5, with χρ = 0.349±0.099.

**Figure 13.**
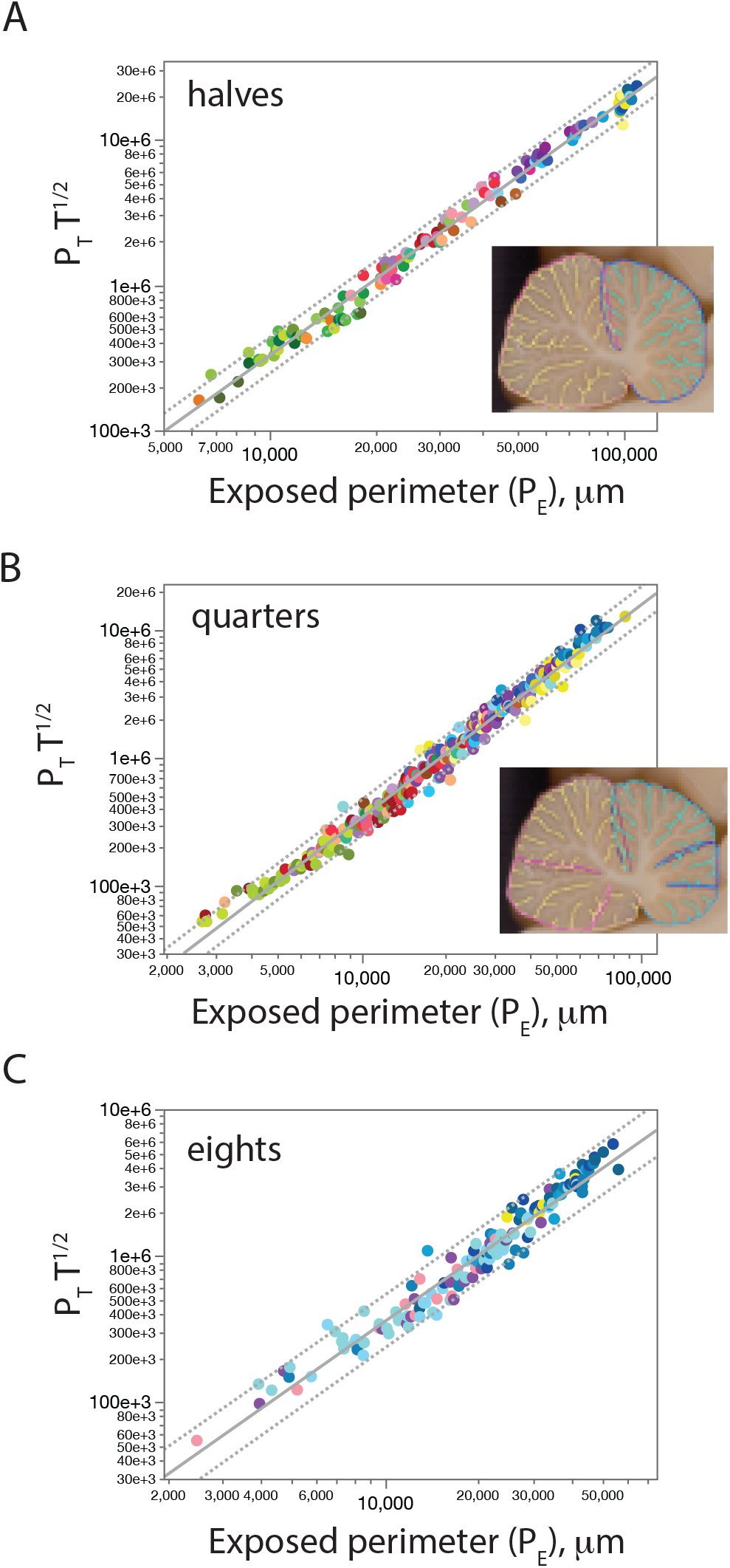
The exponent of the relationship between P_T_ T^1/2^ and P_E_ decreases as progressive levels of analysis exclude the white matter core, which develops prior to expansion of the cerebellar cortical surface in development. **A**, at level 1, which considers P_T_ and P_E_ for the perimeter of each half of the mid-sagittal cerebellar section (the main lobules), defined by the main sulcus, the exponent drops from 1.817 to 1.751±0.016 (r^2^=0.989, p<0.0001). **B**, at level 2, which considers P_T_ and P_E_ for the perimeter of each of the fourths of the midsagittal section of the cerebellum and excludes the core of the subcortical white matter (see inset), the exponent drops even further to 1.652±0.013 (r^2^=0.985, p<0.0001). **C**, at level 3, which considers P_T_ and P_E_ for the perimeter of each of the eights of the two halves of the midsagittal section of the cerebellum, excluding most of the deep cerebellar white matter, the exponent becomes 1.501±0.029 (r^2^=0.956, p<0.0001), identical to the exponent of 1.5 predicted by our model.

The shifting exponents as the white matter core is progressively excluded suggests that the cerebellar cortex as a whole cannot be described as a fractal, but rather as a multi-fractal (Lopes and Beutroni 2009), a geometric object that is also hierarchically composed of substructures of arbitrarily small sizes, but now characterized by a range of scale-dependent fractal dimensions. The findings illustrated in Figure 13 indicate that at the local level, where the contribution of the white matter core is minimal, the cerebellar cortex does fold as a fractal. If this is true, then we should find an invariant fundamental branching element at the appropriate length/thickness ratio derived from our scaling law. We test this prediction in the next section.

### Typical length/thickness ratio of cerebellar folia

Fractal patterns are characterized by repeating units, which in the real world have a minimal size. In the case of the cerebellum, if it does truly fold fractally at the local level (that is, around the white matter core), the ratio p_0_ (defined above) between length of the folia and cortical thickness should be the fundamental structural element of the fractal and should thus be consistent across species and clades, and should be given, as shown in the Methods, 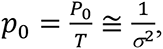 where P_0_ is the average foliar length at the lowest level of branching (which is level 0 for the smallest cerebella, and level 3 for the largest in our sample). Since we find that σ = 0.349±0.099 at branching level 3 (which excludes the bulk of the cerebellar white matter), the predicted length/thickness p_0_ is 8.2±3.8.

Figure 14 shows that this is the case: on branching level 3, across artiodactyl, perissodactyl and carnivoran species, the ratio between the length of folia and the average thickness of the cerebellar cortex is a narrow distribution centered on 7.517±1.809, with 80% of values found between 5.48 and 9.56. The center of the distribution is thus not significantly different from the predicted value of 8.2 for the minimal length/thickness ratio of cerebellar folding, and values are consistent across clades. Importantly, the maximum observed length/thickness ratio is 15.24, just under twice the fundamental unit.

**Figure 14.**
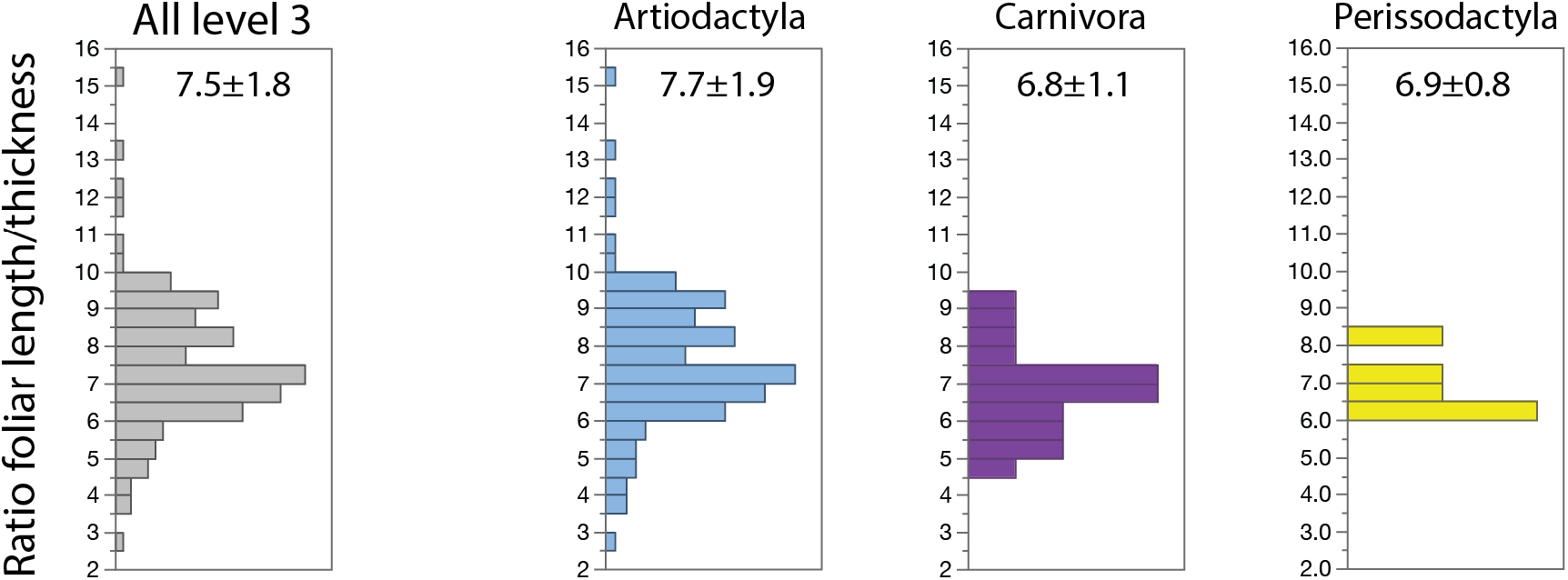
Distribution of the ratio between average foliar length and thickness of the cerebellar cortex at branching level 3 in the mammalian species with the larges cerebella in the dataset is centered around similar values across clades. Values for 8 species with level 3 cerebellar folding are shown in the histogram on the left, and for each clade separately to the right, color-coded as in the other Figures. Mean ratio of foliar length to cerebellar cortical thickness ± standard deviation for each distribution is shown in the respective histograms. At this level of folding, the white matter core has only a minor contribution to the measured perimeter of the cerebellar cortex (from which foliar length is calculated).

Interestingly, the seemingly constant length/thickness ratio of ca. 7.5 is found across species despite clade-specific differences in average thickness of the cerebellar gray matter (Figure 15), with the thinnest cerebellar cortices found in rodents and the thickest in artiodactyls. Although average cerebellar thickness correlates with increasing total perimeter of the mid-sagittal cerebellar pial surface across all species, there is no clear relationship between the total perimeter of a lobule on the midsagittal cerebellar plane and its thickness *within* each clade (Figure 11D); rather, there seems to be a characteristic thickness of the cerebellar cortex to each clade (Figure 15).

**Figure 15.**
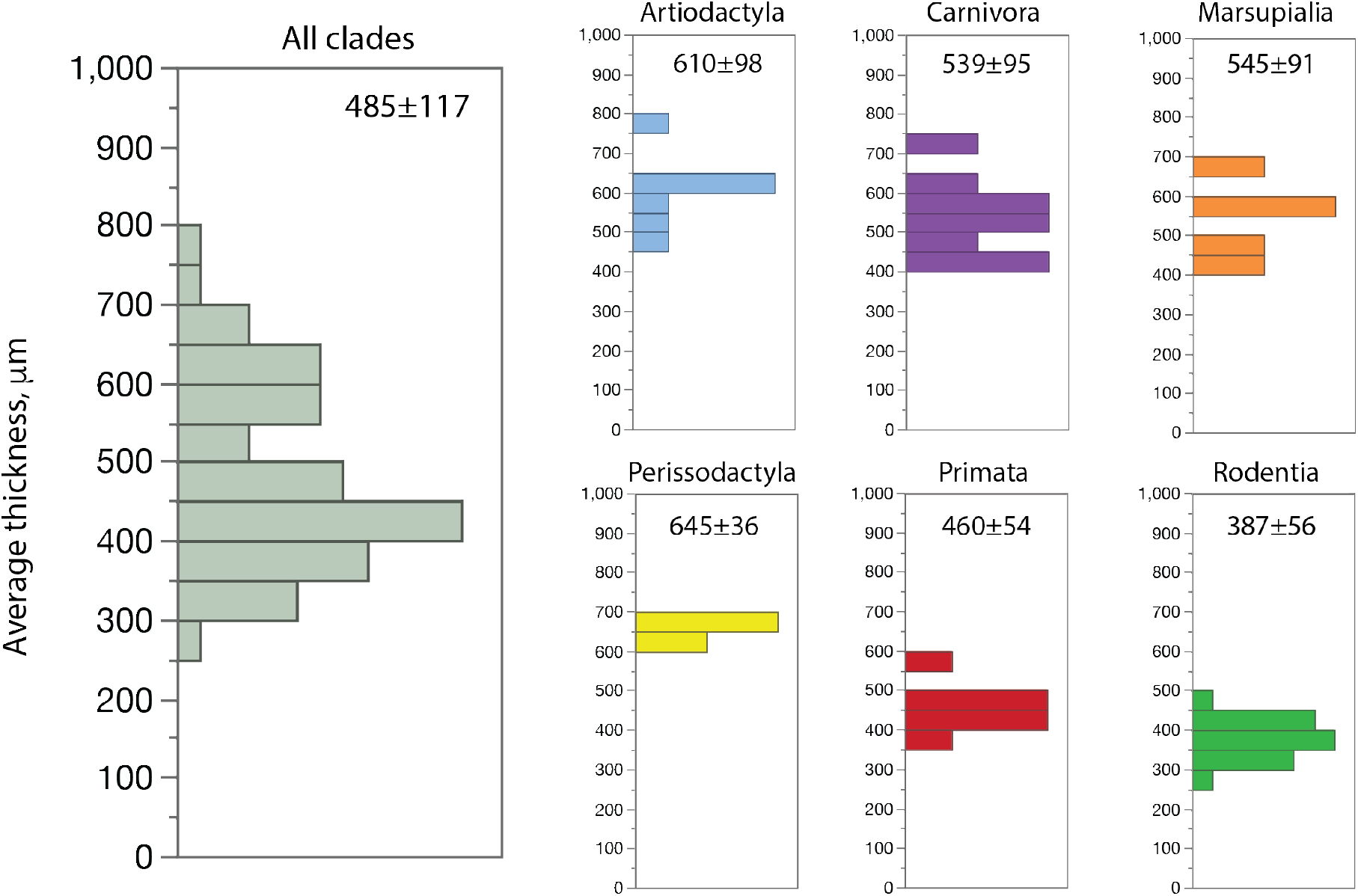
Distribution of average thickness of the cerebellar cortex across the mammalian species in the dataset shows clade-specific values. Values calculated at level 0 for all 53 species in the dataset, ranging from 271 to 800 μm, are shown in the histogram on the left, and for each clade separately to the right, color-coded as in the other Figures. Average cerebellar cortical thickness ± standard deviation for each distribution is shown in the respective histograms.

While significant across clades, the variation in cerebellar cortical thickness is small enough that the typical length/thickness ratio of 7.5±1.8 occurs with a seemingly universal length of cerebellar folia across all clades of approximately 4500 μm (Figure 16), despite the aforementioned variation in degree of folding across clades, as well as the wide range in total perimeter of the cerebella studied. While there is a significant correlation between the perimeter of the mid-sagittal cerebellar pial surface and average foliar length (Figure 11B), the total perimeter of the mid-sagittal pial surface of the cerebellum scales closely and linearly with the number of folia that form it (Figure 12C), suggesting that the small variation in average foliar length does not contribute significantly to cerebellar scaling.

**Figure 16.**
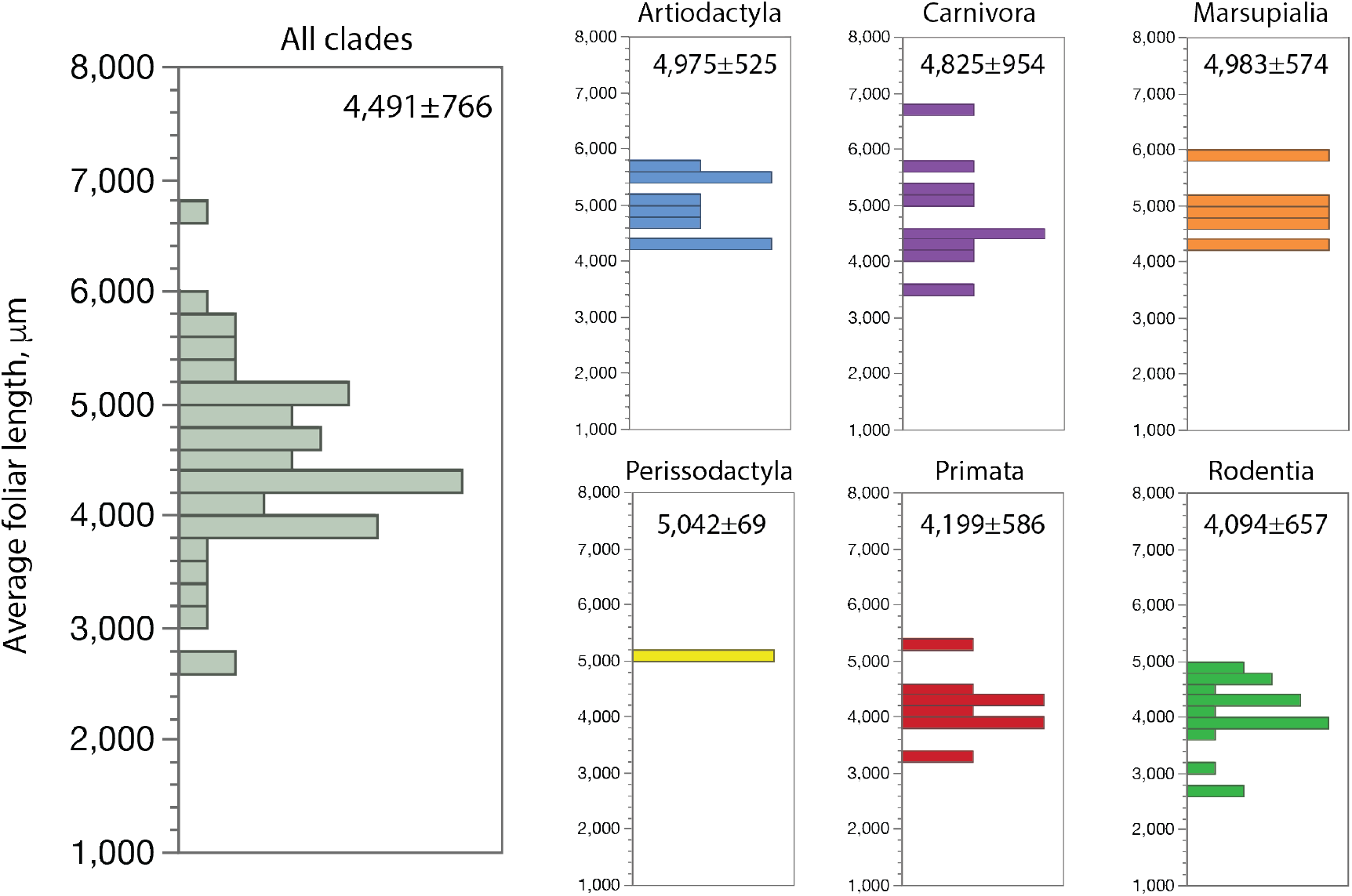
Distribution of average length of cerebellar folia across the mammalian species in the dataset shows consistent lengths of ca. 4.5 mm. Values calculated at level 0 for all 53 species in the dataset, ranging from 2,639 to 6,691 μm, are shown in the histogram on the left, and for each clade separately to the right, color-coded as in the other Figures. Average foliar length ± standard deviation for each distribution is shown in the respective histograms.

We thus conclude that the scaling of the cerebellar surface proportionately to the addition of folia is consistent with a scenario where, once expanding folia reach a critical length/thickness ratio that exceeds the predicted fundamental ratio of 8.2, they fold again, as expected in a fractal process, which, because of the small variation in grey matter thickness, results in a typical foliar length of 4.5 mm.

## DISCUSSION

Here we show, through the alignment of the mid-sagittal pial contours of the cerebellar vermis of different mammalian species, that the spatial arrangement of the main cerebellar folds is clade-specific in pattern. At the same time, the local degree of gyrification of the cerebellar cortex (that is, at the higher branching orders) is almost perfectly predicted by a physical model of the configuration of minimal effective free energy of a volume with a self-avoiding surface that develops under uneven forces (Mota and Herculano-Houzel, 2015), with P_T_ T^1/2^ = k P_E_^1.5^, even if the folding of the cerebellar surface as a whole scales with an exponent that shifts as the contribution of the white matter core is progressively diminished. This indicates that the expanding cerebellar cortex folds locally as a fractal, but the whole cerebellum folds as a multi-fractal, whose fractal dimension is dependent on scale.

We postulate that the multi-fractality and the resulting flow of scaling exponents across folding levels and deviation from the expected value are due to the precedence of the white matter core that forms in the cerebellum before the cerebellar grey matter surface begins to expand in development. Our gyrification model applies to a self-avoiding sheet of grey matter that connects through and expands together with the underlying white matter, so that their volumes are tightly correlated (Mota et al., 2019). Cerebellar development differs markedly in this aspect from cerebral cortical development in that the core of the cerebellar white matter forms early and well before the external granule cell layer expands and gives rise to the layered cerebellar cortical grey matter. As a consequence, this pre-existing cerebellar white matter contributes to the measured section perimeter of the mid-sagittal section without fully contributing to the gyrification dynamics, resulting in a scaling exponent that deviates from the predicted value. Once the white matter core is no longer a significant contributor to the cerebellar surface, the midsagittal section of the cerebellar gray matter does scale as a fractal, similar to the cerebral cortex.

We thus conclude that a similar mechanism driven primarily by axonal elongation dynamics and self-avoidance of surfaces (Mota and Herculano-Houzel, 2015; Wang et al., 2022) generates gyrification of the cerebellar and cerebral cortices, as can be expected from physics alone, despite the independent developmental origins of these two structures and their disparate developmental programs, cytoarchitecture and connectivity. In the case of the cerebellum, such folding is locally well-approximated by a fractal of dimension d_f_ = 1.5, with a typical terminal length/thickness ratio of ∼8.2. A fractal mechanism of folding explains how come essentially no cerebellar folia have a length/thickness ratio greater than twice this value: we expect that, during development, any stretch of cerebellar cortical surface whose perimeter on the sagittal plane exceeds 8.2 times its thickness will tend to fold in two, such that no folium is larger than twice that ratio.

Our findings have several important consequences for the understanding of vertebrate brain evolution. First, the universality of the relationship that describes the degree of folding of both the cerebellar and cerebral cortices according to the minimization of effective free energy of their grey matter volumes indicates that rather than being a biological property that evolved specifically in some clades but not others, cortical folding is driven by a universal mechanism based purely upon physical determinants. It is neither a direct consequence of increasing numbers of neurons nor a requirement for increasing number of neurons, nor is it directly related to brain size or cortical volume, and probably not to any other biological determinant. As the physical model was found to apply in the same manner to crumpled sheets of paper as with mammalian cortices, we propose that folding is “an intrinsic, fractal property of a self-avoiding surface, whether biological or not, subjected to crumpling forces,” be that surface a sheet of paper or biological tissue (Mota and Herculano-Houzel, 2015). In this case, we can thus explain the much more pronounced folding of the cerebellar surface compared to that of the cerebral cortical surface in any species as largely due to the larger surface area and thinner thickness of the former, and not to any intrinsic, genetic, or evolutionary differences in the fundamental nature of each cortex.

Secondly, our findings explain the apparent clade-specificity of the degree of cerebellar folding across bird clades described by Iwaniuk et al. (2006) as a consequence of using cortical perimeter, a non-universal predictor of folding, as independent variable (Iwaniuk et al., 2006), just as one could describe the degree of cerebellar folding as clade-specific functions of the numbers of cerebellar neurons as shown here (Figure 12B). As we show here, the universal predictor of the degree of gyrification is the ratio between cortical perimeter and the square root of cortical thickness in two-dimensional brain sections (which is proportional to the ratio between cortical surface area and thickness in three-dimensional structures) – that is, both cortical surface extension and thickness are key determinants of the degree of folding. We predict that extending our analysis to birds will show that our model also explains universally the degree of folding of the bird cerebellum.

Thirdly, our finding that there are distinct, clade-specific patterns to the spatial arrangement of the main cerebellar lobes, lobules and folia directly contradicts the previously held assumption that all mammalian cerebella follow a common morphological plan (Stroud, 1895; Sultan and Braitenberg, 1993; Sultan & Glickstein, 2007; Voogd & Glickstein, 1998). We believe that our findings cast sufficient doubt on a shared morphological plan that attempts to squeeze the cerebella of different species into the ten-lobule paradigm will from now on be considered shoehorning, for the clade-specific patterns revealed here imply that we cannot treat all mammalian cerebella as functionally equivalent in their correspondence between anatomical landmarks (sulci and lobules) and functional divisions. More specifically, the human cerebellum, as a primate cerebellum, cannot be compared to a scaled-up version of a mouse (rodent) cerebellum. Because this difference in the spatial placement of the main sulci may impact (or reflect) the location of functional zone landmarks in the cerebella of different mammalian species, our demonstration of clade-specific spatial patterns of cerebellar folding indicate that we need systematic comparative studies across species within a clade, and then across clades, to understand the correspondence between the location of cerebellar sulci and functional specialization in the cerebellum. Only then will we be able to consider the extent to which findings from research with animal models can be generalized from one species to another. Importantly, while white matter connectivity may play a role in the placement of main sulci in the mammalian cerebral cortex (Van Essen, 1997), it is unlikely that white matter connectivity plays a role in establishing clade-specific patterns in the cerebellum, given the lack of horizontal connections through the cerebellar white matter (Voogd and Glickstein, 1998; Bush and Allman, 2003). Additionally, it was recently shown that the cerebellar cortex folds initiates with differential expansion of the outer layer, without an obvious cellular pre-pattern, and with differential expansion of lobules that correlates with the progressive subfolding of the initial folds (Lawton et al., 2019), which is consistent with the fractal scaling of cerebellar folding that we propose here. We thus find it more likely that the placement of cerebellar folds is controlled by clade-specific genetic factors that may lead to differential patterns of cortical expansion and/or placement of anchoring centers (Sudarov and Joyner, 2007) that might accompany evolutionary divergence between the mammalian clades.

These considerations about the placing of the main cerebellar folds leads to the fourth evolutionary implication of our findings: if the *mechanism* of cerebellar folding is not an evolved feature, but rather a consequence of fundamental physics, with hierarchical placement of successive folds once a critical length is reached, it becomes unlikely that there is some sort of an “ancestral cerebellar pattern”. Our results reinforce Mota and Herculano-Houzel’s proposal in 2015 that, given a set surface area and thickness, subjected to uneven forces, a sheet of grey matter will always produce the same degree of folding regardless of the mammalian clade to which it belongs, or the number of neurons it contains.

The fractal pattern of cerebellar folding revealed by our present findings begs a developmental investigation of several predictions that we can make. While our qualitative analysis was restricted to the visual analysis of already formed adult cerebella, their overlaid contours share a pattern suggestive of a hierarchical order of folding, with the specific prediction that the deepest main fissure forms first, and that the formation of the subsequent folds follows in the same order as their apparent depth in the adult cerebella. Thus, we predict that the two horizontal folds which divide the cerebellum from two halves into quarters (so to speak) are next to form after the two halves formed by the main fissure expand to the critical foliar length, given that they have the highest level of coincidence across species. As the hierarchical order of folding continues, the decreased coincidence in sulcal placement most likely arises as a result of certain lobes or lobules expanding at a faster rate than others, as suggested by Lawton et al. (2019). Thus, an uneven growth of lobules that is shared in the same manner between species of the same clade could be the mechanistic reason behind the formation of the clade-specific spatial arrangement patterns. Developmental studies would make it possible to observe empirically the order in which foliation occurs, as well as how the spatial placement of the main lobes arises as the cerebellum expands. Comparative studies of cerebellar development would thus reveal how clade-specific patterns of folding arise, even as folding itself occurs predictably due to the physical properties of biological tissue.

Finally, a lingering question at this point is whether the remarkable degree of folding of even very large cerebella, such as those of humans, elephants, and cetaceans, as well as in the minute cerebella of bats, also follows the universal relationship described here, while maintaining the spatial pattern that is typical of their clades. These studies, which will test the validity of our model not only in humans but also in a number of species known to have relatively larger cerebella (Maseko et al., 2012; Clark et al., 2001), whether because the number of neurons in the cerebellum is extraordinarily large (Herculano-Houzel et al., 2014) or because the cerebral cortex is relatively small in volume (Herculano-Houzel et al., 2020), are currently underway.

## Acknowledgments

Supported by the James S. McDonnell Foundation (SHH), Fundação Serrapilheira (grant Serra-1709-16981) and CNPq (PQ 2017 312837/2017-8) (BM), and Vanderbilt University startup funds (SHH).

## Author contributions

ARY performed research, analyzed data and wrote the paper; CCS, PRM and JHK contributed research materials; BM designed research, analyzed data, and wrote the paper; SHH designed research, performed research, analyzed data and wrote the paper.

